# Astrocytes mediate the pro-cognitive value of α7nAChRs and of α7nAChR-targeting therapeutics

**DOI:** 10.64898/2026.04.16.719027

**Authors:** Yifan Wu, Michaela Tolman, Yanchao Dai, Sarah Walsh, Humza Agha, Katheryn B. Lefton, Hannah An, Rachel Manno, Philip G. Haydon, Thomas Papouin

**Affiliations:** Washington University in St Louis, School of Medicine, Department of Neuroscience, St Louis, MO USA; Tufts University School of Medicine, Neuroscience Department, Boston, USA

## Abstract

The α7-nicotinic acetylcholine receptor (α7nAChR) has driven extensive research over the past three decades for its pro-cognitive potential. It is the leading druggable target for the cognitive deficits associated with schizophrenia and has motivated major pharmaceutical and clinical efforts to ameliorate similar impairments in other neurological disorders, such as Alzheimer’s disease (AD). Yet, a systematic evaluation of the role played by α7nAChR in cognition, and its mechanistic underpinnings, is still lacking. Here we report that α7nAChRs on principal and inhibitory forebrain neurons are largely inconsequential to mouse behavior, including in domains that are most sensitive to schizophrenia-related cognitive impairments. By contrast, loss of α7nAChR from astrocytes produces profound behavioral alterations that are cognitive domain-specific, are time-of-day dependent, coincide with reduced levels of the N-methyl D-aspartate receptor (NMDAR) co-agonist D-serine, and are fully restored by D-serine supplementation. Further, an α7nAChR partial agonist previously evaluated in Phase III trials for cognitive enhancement in schizophrenia and AD fails to augment behavior in mice lacking astrocytic α7nAChRs. Together, these findings identify astrocytes and D-serine/NMDAR signaling as a central mechanism through which α7nAChR, a major drug target, promotes cognitive behavior.

## Main

Cognitive impairments are highly prevalent in patients suffering from schizophrenia, including learning deficits and reduced social functioning^1^. These behavioral impairments precede the onset of positive symptoms, and their severity is predictive of patient’s functional outcomes (e.g., occupational integration). However, the cognitive symptoms of the disease remain largely insensitive to first and second-generation antipsychotic medications, and no treatments are currently available to alleviate them^1^. In response, the US National Institute of Mental Health launched the Measurement and Treatment Research to Improve Cognition in Schizophrenia (MATRICS) initiative^2^, with the mandate to establish a translational framework for improving the discovery and testing of cognition-enhancing treatments. In its effort, the MATRICS consensus identified the α7 nicotinic acetylcholine receptor (α7nAChR) as the leading druggable target to alleviate the cognitive deficits of schizophrenia^2^. This ranking reflected evidence of reduced α7nAChR immunoreactivity in brain tissue of patients^3,4^, and human genetic associations linking *CHRNA7* variants, *CHRNA7* duplications, and the chromosomal region containing the *CHRNA7* locus to cognitive impairments typical in schizophrenia^5,6^. It also echoed pharmaco-behavioral evidence of the pro-cognitive properties of α7nAChRs in rodents^7^. Pharmaceutical efforts have since produced myriad α7nAChR agonists and positive allosteric modulators of which 12 were tested in over 30 unique clinical trials for their safety and efficacy at improving cognition in schizophrenia, as well as in Parkinson’s disease (PD) and Alzheimer’s disease (AD) where similar cognitive impairments are a major clinical feature^8–10^. Yet, a direct and systematic assessment of the contribution of α7nAChR to cognitive behavior, and candidate mechanisms linking its putative pro-cognitive value to specific cellular processes, are still lacking.

Owing to their high calcium permeability^11,12^, it is generally accepted that α7nAChRs regulate Ca^2+^-dependent neuronal excitability, synaptic transmission, and synaptic plasticity in rodents and primates^13–16^, which is thought to underly their participation in cognitive processes such as learning^16^. Despite predominant focus on α7nAChR functions on various populations of neurons, these receptors are ubiquitously expressed in the brain, including by non-neuronal cells^17^. Since acetylcholine signals diffusively via volume transmission to engage receptors on many cell-types and sub-cellular locations, rather than on defined post-synaptic neuronal targets^18^, the participation of non-neuronal cells such as astrocytes to α7nAChR-dependent behaviors is potentially relevant. Here, we thus set out to systematically evaluate the importance of α7nAChRs expressed by forebrain principal neurons, inhibitory neurons, and astrocytes, to mouse behavior as a step towards elucidating cellular and molecular underpinnings of α7nAChR relation to cognition. We generated three knockout (KO) mouse lines with equivalent penetrance, high specificity for the targeted cell-types, and conditional ablation in adulthood, and examined their performance across behavioral domains.

## α7nAChR deletion from excitatory neurons is largely inconsequential to behavior

We first set out to investigate the behavioral relevance of α7nAChR expressed on excitatory neurons, considering that the bulk of *Chrna7* expression in the brain occurs in these cells^19^. We used *Chrna7^fl/fl^* mice in which exon 4 that codes the N-terminal agonist-binding domain of α7nAChR can be deleted in a Cre-dependent manner^20^ and crossed them with Camk2a::CreER^T2^ mice to create inducible excitatory neuron-specific α7nAChR KOs (eN-α7^KO^, Fig.1A). The Cre-driver line was chosen for its documented cell-type specificity, efficiency, and CreER^T2^ fusion protein, allowing targeted recombination via tamoxifen treatment in adulthood^21,22^, which we validated using four complementary approaches (Fig.1B-D and Fig.S1A-D). First, we crossed these mice with the Ai9 reporter line (LSL-tdTomato) and found that 93-99% of Cre-expressing cells (tdTomato-positive) were neurons (NeuN-positive) across five brain regions quantitatively examined, with 49-59% of all neurons recombined (Fig.1B, and Fig.S1A). Further, *Chrna7* mRNA and native genomic DNA levels were reduced by ∼40% in eN-α7^KO^ mice (Fig.1C, D), with clear evidence of recombined alleles on PCR gels, together confirming genomic recombination (Fig.1D). Finally, 2-photon Ca²⁺ imaging (2-PLSM, Fig.S1B) showed that α7nAChR-mediated responses to brief applications of choline, a preferred agonist^23^, were abolished in Cre-positive neurons but preserved in Cre-negative neurons, confirming loss of receptor function (Fig.S1C, D).

**Figure 1:**
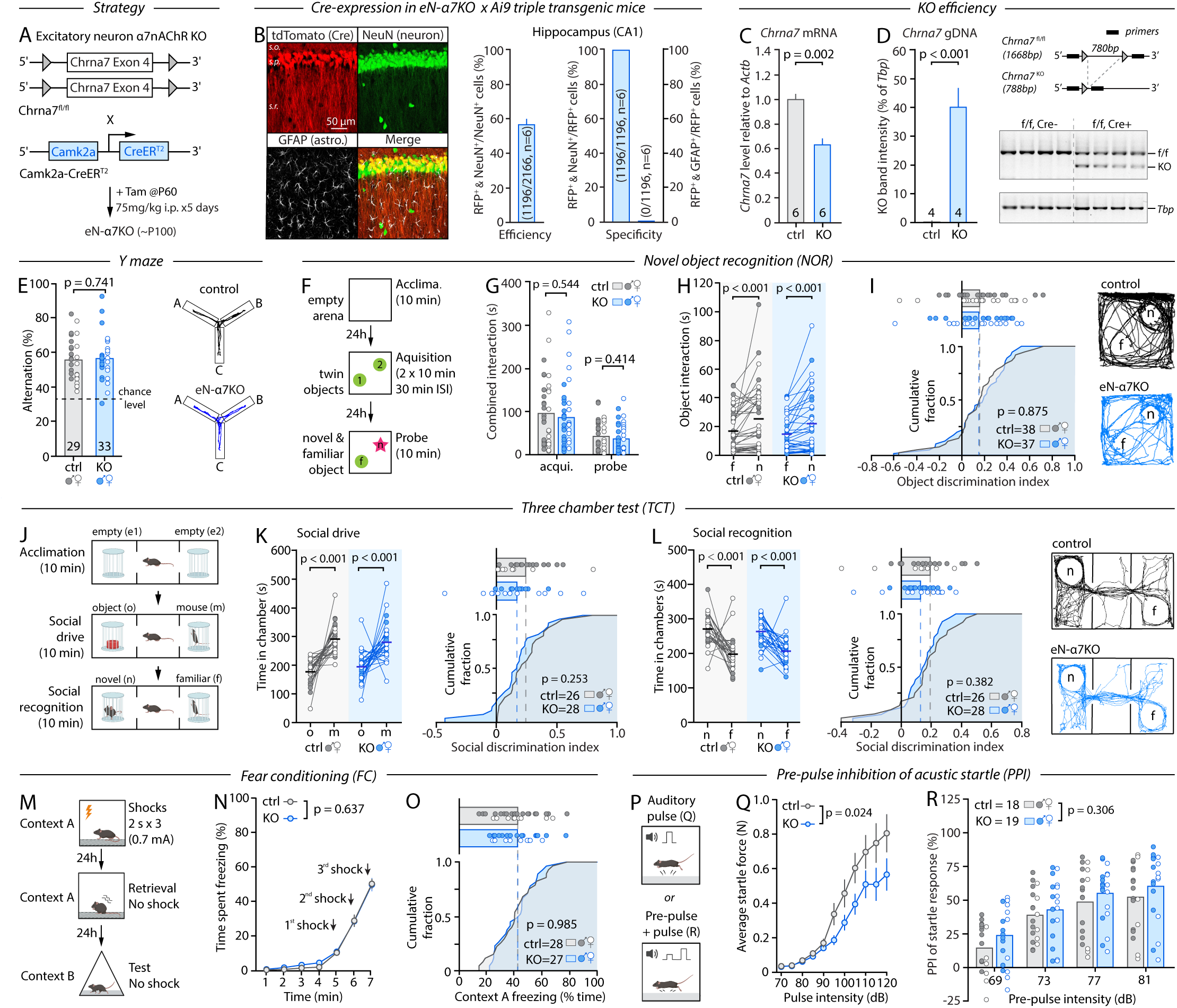
Loss of α7nAChR from excitatory neurons is inconsequential to MCCB-related behavior. **A**, Genetic strategy and timeline for eN-a7^KO^. **B**, IHC and analysis of recombination specificity and efficiency in eN-α7^KO^ crossed with the Ai9 reporter line. **C**, RT-qPCR analysis of *Chrna7* transcript levels in brains of control and eN-α7^KO^ mice. **D**, *Left*, PCR of genomic DNA in control and eN-α7^KO^ mice. *Right*, schematic of predicted amplicon size in control and eN-a7^KO^ mice with corresponding gel images. **E**, Percentage of spontaneous arm alternations in the Y-maze and representative mouse trajectories. **F**, Schematic of the NOR paradigm. **G**, Time spent exploring both objects during the acquisition and probe phase. **H**, Time spent interacting with the novel (n) and familiar (f) object during the probe phase of the NOR. **I**, Computed object discrimination index of control and eN-α7^KO^ mice shown as cumulative fraction (bottom) and individual data (top), and representative mouse trajectories during the probe phase. **J**, Schematic of the TCT paradigm. **K**, *Left*: Time spent in the chamber containing the object (o) and the conspecific (m) during the second phase of the TCT. *Right*: Computed social discrimination index as shown in I. **L**, *Left*: Time spent in the chamber containing the familiar (f) and novel mouse (n) during the third phase of the TCT. *Middle*: Computed social discrimination as shown in I and K. *Right*: Representative mouse trajectories. **M**, Schematic of the contextual FC paradigm. **N**, Percentage of time spent freezing in response to shocks during the fear learning phase of the FC. **O**, Percentage of time spent freezing during the fear retrieval phase of the FC shown as cumulative fraction as in I, K, and L. **P**, Schematic of the auditory startle and PPI paradigms. **Q**, Startle responses to auditory pulses of increasing intensity. **R**, PPI of the auditory startle response for 4 pre-pulse intensities. Bar graphs and thick horizontal bars in pair-wise graphs show averages, here and thereafter. Filled circles show males, open circles show females, here and thereafter. Unpaired Student’s *t*-test (D, E, I, O), Mann-Whitney test (C, G, K, L), paired Student’s *t*-test (K), Wilcoxon test (H, L), two-way repeated measures ANOVA with Sidak’s multiple comparisons test (N, R) and mixed-effects analysis (Q) were used.

Given the longstanding relevance of α7nAChR to the cognitive aspects of schizophrenia, we first assessed how their loss in excitatory neurons altered cognitive function. We specifically focused on behavioral domains encompassed by the MATRICS Consensus Cognitive Battery (MCCB)^2^. The MCCB is a benchmark developed by the MATRICS consensus to evaluate cognitive abilities in schizophrenia trials. It was rapidly adopted as a primary cognitive endpoint by the US Food and Drug Administration (FDA) and non-US regulatory agencies, and has become an international reference routinely used in clinical trials worldwide^24,25^. The MCCB encompasses seven major cognitive domains (working memory, attention/vigilance, verbal/semantic learning, visual/spatial learning, social cognition, processing speed/sensory gating, and reasoning, Marder, 2006) that can be evaluated in mice using well-established behavior tests^26–30^, making it a logical framework to assess eN-α7^KO^ mice. We excluded reasoning from our assessment owing to its contentious nature in mice. Since mouse behavioral assays rarely map cleanly onto discrete behavior domains, we used a battery of complementary tasks to capture MCCB domains collectively: the spontaneous Y-maze, novel object recognition (NOR), three-chamber test of social drive and social memory (TCT), contextual fear conditioning (FC), and pre-pulse inhibition (PPI) of auditory startle responses. Owing to the state-dependent nature of cholinergic signaling^31^, mice were tested in their subjective day (Zeitgeber time 0, ZT0) when they were still awake and active.

Male and female eN-α7^KO^ mice appeared healthy, weighed normally, were indistinguishable from their same-sex tamoxifen-treated littermate controls by blinded experimenters when handled or observed in their home-cage and, surprisingly, showed no phenotype in all but one metric across the entire battery of tests (Fig.1E-R). Indeed, eN-α7^KO^ mice showed nominal rates of spontaneous arm alternations in the Y-maze (Fig.1E, and Fig.S1E), performed as well as their control littermates at the NOR task by all metrics (Fig.1F-I and Fig.S1F), showed expected social drive and social memory in the TCT social assay (Fig1J-L. and Fig.S1G,H), displayed normal acquisition of fear response to shocks and normal freezing upon retrieval in the same context 24 hours later (Fig.1M-O and Fig.S1I), and had similar PPI of acoustic startle responses across a set of 4 pre-pulse intensities compared to controls (Fig.1P,R) besides a marked ∼30% reduction in the magnitude of their startle responses to auditory stimuli (Fig.1Q). Mouse behavior apparatus/tests inherently engage a composite of behavioral functions. For instance, completing the NOR involves ambulation, attention, tactile discrimination, visual processing and, even under low lighting, can evoke anxiety-like responses. Normal performance in the above assays thus suggested that the lack of eN-α7^KO^ phenotype extended to non-MCCB behaviors. To test this directly, we expanded our battery by including measures of locomotion (in open field, OF), anxiety-like (elevated plus maze, EPM), pleasure-seeking (two-bottle sucrose preference test, SPT), and nesting behavior (Fig.S3A-F), based on reports that linked α7nAChRs to these behaviors and owing to their relevance to aspects of schizophrenia such as avolition, anhedonia and anxiety^10^. Shockingly, eN-α7^KO^ mice were indistinguishable from their littermate controls in these assays as well (Fig.S3A-F), portraying a general lack of phenotype in these mice across a total of 37 different metrics in 12 behavior paradigms.

## Ablation of α7nAChRs from inhibitory neurons has minimal impact on MCCB behaviors

Recent reports suggest that the constitutive deletion of α7nAChRs from inhibitory neurons from birth yields overlapping behavioral impairments with those reported in global KOs^32^. Therefore, we next generated conditional inhibitory neuron-specific α7nAChR KO mice (iN-α7^KO^), by crossing *Chrna7^fl/fl^* mice with the Gad2::CreER^T2^ driver line (Fig.2A). Systematic examination confirmed the effective and cell-specific recombination of *Chrna7* in iN-α7^KO^ mice, as for eN-α7^KO^ mice, with comparable penetrance and regionality (Fig.2B-D and Fig.S2A). Like eN-α7^KO^, iN-α7^KO^ males and females appeared healthy, displayed normal behavior when observed in their home-cage or upon handling, and were undistinguishable from their same-sex littermate controls by blinded experimenters (Fig.S4A,B). Strikingly, when evaluated in the same battery of tests under the same conditions, they too showed no overt change in behavioral performance. iN-α7^KO^ mice performed nominally in the Y-maze (Fig.2E and Fig.S2B) and the NOR (Fig.2F-I and Fig.S2C), had expected social preference and social recognition in the TCT (Fig.2J-L and Fig.S2D,E), and showed normal auditory startle responses and PPI (Fig.2P-R). Notably, iN-α7^KO^ mice displayed a mild reduction in context-specific fear retrieval (Fig.2M-O, but see Fig.S2F), indicative of potential impairments in contextual fear learning and memory, consistent with the proposed role of interneuron α7nAChRs in hippocampal theta oscillations and learning^33^. Contrary to the deficits observed in mice lacking α7nAChRs from interneurons from birth^33,34^, we did not observe deficits in social recognition or Y-maze alternations, suggesting that these impairments may have been developmentally acquired.

**Figure 2:**
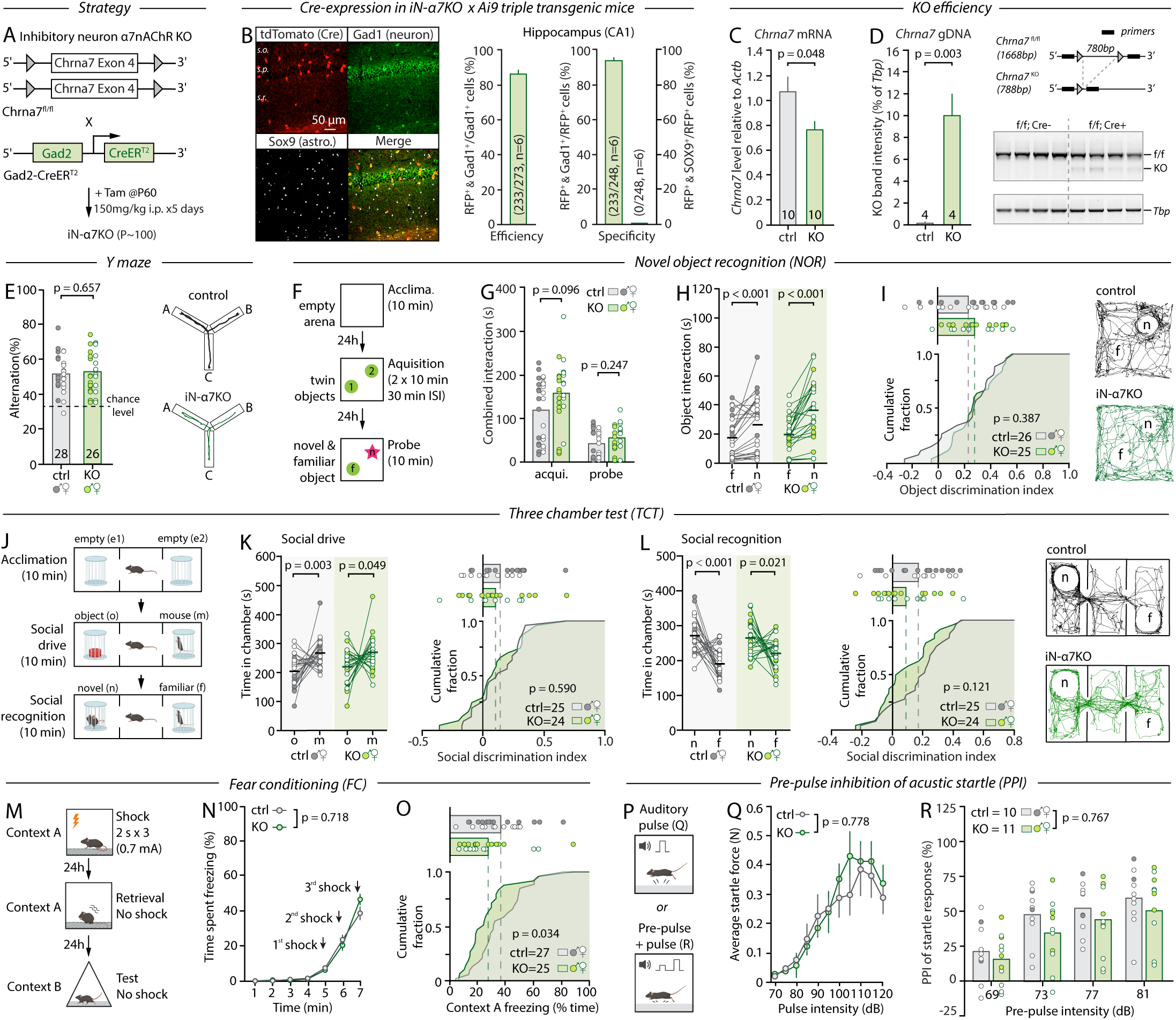
Loss of α7nAChR from inhibitory neurons has little impact on MCCB behavior. **A**, Genetic strategy and timeline for iN-a7^KO^. **B**, IHC and analysis of recombination specificity and efficiency in iN-α7^KO^ crossed with the Ai9 reporter line. **C**, RT-qPCR analysis of *Chrna7* transcript levels in brains of control and iN-α7^KO^ mice. **D**, *Left*, PCR of genomic DNA in control and iN-α7^KO^ mice. *Right*, schematic of predicted amplicon size in control and iN-a7^KO^ mice with corresponding gel images. **E**, Percentage of spontaneous arm alternations in the Y-maze and representative mouse trajectories. **F**, Schematic of the NOR paradigm. **G**, Time spent exploring both objects during the acquisition and probe phase. **H**, Time spent interacting with the novel (n) and familiar (f) object during the probe phase of the NOR. **I**, Computed object discrimination index of control and iN-α7^KO^ mice shown as cumulative fraction (bottom) and individual data (top), and representative mouse trajectories during the probe phase. **J**, Schematic of the TCT paradigm. **K**, *Left*: Time spent in the chamber containing the object (o) and the conspecific (m) during the second phase of the TCT. *Right*: Computed social discrimination index as shown in I. **L**, *Left*: Time spent in the chamber containing the familiar (f) and novel mouse (n) during the third phase of the TCT. *Middle*: Computed social discrimination as shown in I and K. *Right*: Representative trajectories. **M**, Schematic of the contextual FC paradigm. **N**, Percentage of time spent freezing in response to shocks during the fear learning phase of the FC. **O**, Percentage of time spent freezing during the fear retrieval phase shown as cumulative fraction as in I, K, and L. **P**, Schematic of the auditory startle and PPI paradigms. **Q**, Startle responses to auditory pulses of increasing intensity. **R**, PPI of the auditory startle response for 4 pre-pulse intensities. Unpaired Student’s *t*-test (C, D, E, I, K right, L right), Mann-Whitney test (G, O), paired Student’s *t*-test (H, K left, L left), Wilcoxon test (H, K) and two-way repeated measures ANOVA with Sidak’s multiple comparisons tests (N, Q, R) were used.

Interestingly, probing iN-α7^KO^ mice in non-MCCB domains, we found several behavioral differences (Fig.S4B-F). Both male and female KOs displayed markedly improved nesting scores (Fig.S4F), increased sucrose preference (Fig.S4C), and a significant reduction in open-arm poking in the EPM besides no other indications of increased anxiety (Fig.S4C,D). Altogether, these findings suggest that while α7nAChRs on inhibitory neurons are relevant to mouse behavior, the deletion of α7nAChRs from inhibitory or excitatory neurons is largely inconsequential to MCCB-related functions.

## Astrocytic α7nAChRs drive transmembrane calcium dynamics

Astrocytes are now recognized as ubiquitous orchestrators of brain circuits that actively shape behavior^35–37^. Growing evidence suggests that astrocytes mediate some of the circuit effects of neuromodulators across species^20,38–44^, including those of acetylcholine^38–41^. However, single cell and single nuclei RNAseq datasets show very low, or sometimes undetectable levels of *Chrna7* in astrocytes^45,46^. Since α7nAChRs are highly Ca^2+^-permeable ligand-gated ion channels, we verified the presence of functional α7nAChRs at the surface of astrocytes by performing 2-PLSM imaging studies of Ca^2+^ dynamics in acute hippocampal slices (Fig.S5A). We found that bath application of the α7nAChR-specific agonist PNU-282987 (500 nM), in the presence of tetrodotoxin (TTX, 1 µM) to silence neuronal activity, did not evoke whole cell responses in lck-GCaMP6f-expressing astrocytes but significantly impacted Ca^2+^ activity in microdomains (Fig.S5B-E). PNU-282987 increased the amplitude and the area under the curve (AUC) of spontaneous Ca^2+^ transients most notably, along with changes in kinetics, all of which were abolished in the presence of the α7nAChR-specific antagonist methyllycaconitine (MLA, 100-500 nM, Fig.S5B). α7nAChRs are ionotropic but were also proposed to signal through non-canonical metabotropic pathways, including Gαq signaling^47^. Seeing how some astrocyte Ca^2+^ activities rely on Gαq pathways, we tested the role of this signaling modality. We first found that the effect of the α7nAChR agonist PNU-282987 was enhanced in the presence of a positive allosteric modulator (PNU-120596, 2 µM) that increases agonist-evoked peak currents through α7nAChR channel by delaying channel desensitization ^12^. This suggested a reliance on transmembrane Ca^2+^ fluxes (Fig.S5B). Consistently, the effect of the agonist PNU-282987 was not affected by manipulations that deplete intracellular Ca^2+^ stores in astrocytes (thapsigargin, 1 µM), but was attenuated by omitting extracellular Ca^2+^, and fully abolished by the further addition of extracellular Ca^2+^ chelator (EGTA, 2 mM, Fig.S5B). Altogether, these results confirm the expression of functional α7nAChRs on astrocytes that influence microdomain Ca^2+^ dynamics via local transmembrane Ca^2+^ influx.

## Loss of astrocytic α7nAChRs drives MCCB-specific behavioral deficits

Based on the above results, we next examined the potential relevance of astrocytic α7nAChRs to MCCB-related behaviors. We used the Glast::CreER^T2^ line to drive *Chrna7* recombination in astrocytes and generate astrocyte-specific α7nAChR KO mice (A-α7^KO^, Fig.3A). This Cre-driver line was favored over the more popular Aldh1l1::CreER^T2^ due to high and widespread peripheral expression of Cre in the latter ^48^. In addition, the recombination penetrance reported for Glast::CreER^T2^, although lower, compares more closely to that achieved with Camk2a::CreER^T2^ and Gad2::CreER^T2^ for targeted cell-types^21,22^ allowing for a more direct comparison. Triple-transgenic crossing with Ai9 reporter mice confirmed widespread astrocyte-specific recombination with a penetrance reminiscent of that achieved in eN-α7^KO^ and iN-α7^KO^ mice (Fig.3B and Fig.S6A). Importantly, as previously reported^46,49^, we found that Cre was also sparsely expressed in a subset of adult new-born neurons in the dentate gyrus (12% of all DG neurons) where the astrocyte specificity of Cre-recombination was suboptimal (Fig.S6A), which we address below. *Chrna7* mRNA levels were reduced by 22 ± 8% in A-α7KO mice (Fig.3C), and recombinant bands were detected by PCR (Fig.3D). Further, the amplitude and AUC of spontaneous Ca^2+^ transients were attenuated in astrocytes of A-α7^KO^ mice compared to littermate controls (Fig.S5B, H) and unresponsive to the α7nAChR agonist PNU-282987 (Fig.S5B,C,F,G). Together, these observations confirm the effective and astrocyte-specific deletion of α7nAChR in A-α7^KO^ mice suitable for behavioral studies, with a penetrance of recombination directly comparable to that of eN-α7^KO^ and iN-α7^KO^ mice.

**Figure 3:**
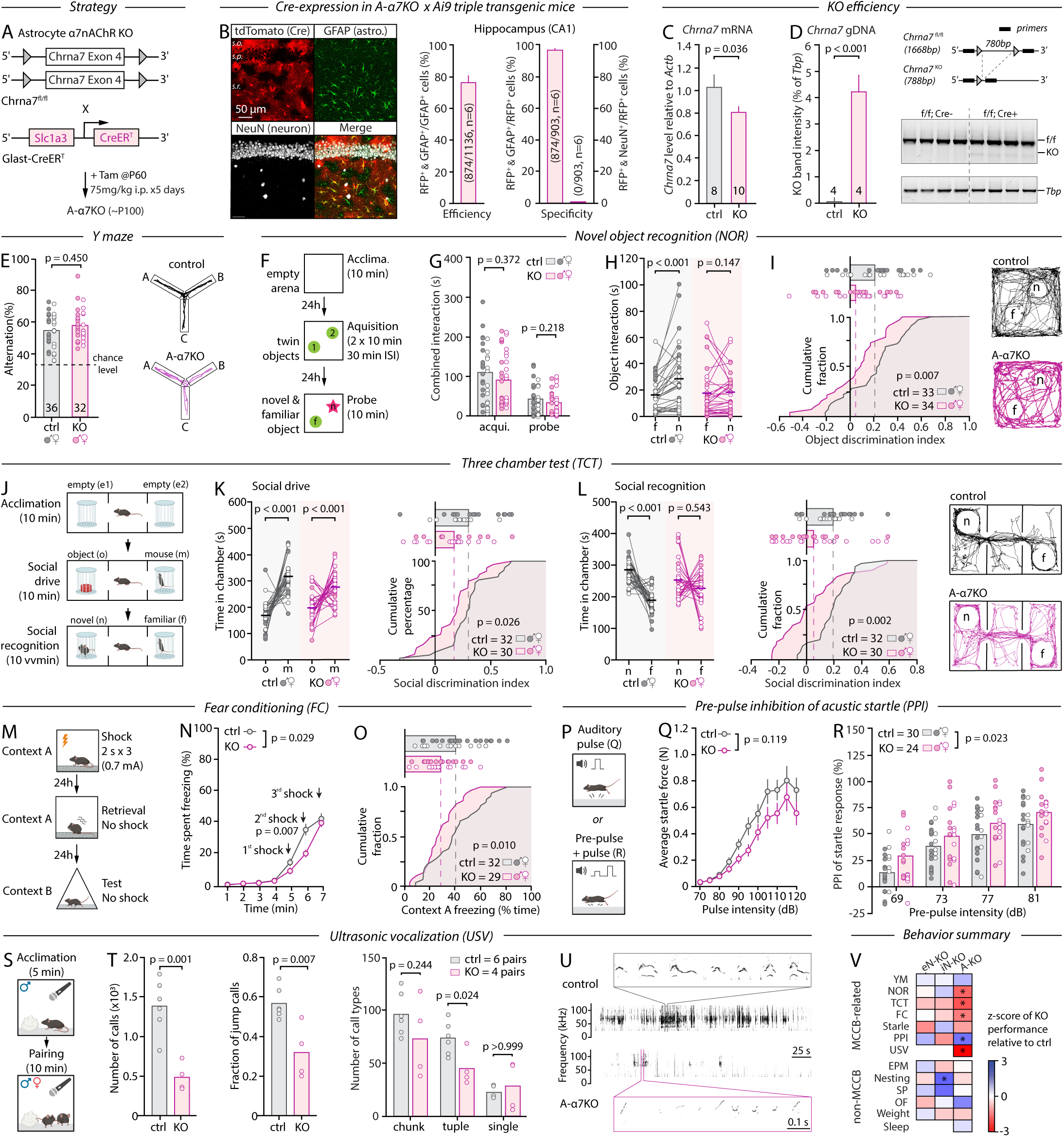
Mice lacking α7nAChR from astrocytes have profound behavior alterations in MCCB domains. **A**, Genetic strategy and timeline for A- α7^KO^. **B**, IHC and analysis of recombination specificity and efficiency in A-α7^KO^ crossed with the Ai9 reporter line. **C**, RT-qPCR analysis of *Chrna7* transcript levels in brains of control and A-α7^KO^ mice. **D**, *Left*, PCR of genomic DNA in control and A-α7^KO^ mice. *Right*, schematic of predicted amplicon size in control and A-α7^KO^ mice with corresponding gel image. **E**, Percentage of spontaneous arm alternations in the Y-maze and representative mouse trajectories. **F**, Schematic of the NOR paradigm. **G**, Time spent exploring both objects during the acquisition and probe phase. **H**, Time spent interacting with the novel (n) and familiar (f) object during the probe phase of the NOR. **I**, Computed object discrimination index of control and A-α7^KO^ mice shown as cumulative fraction (bottom) and individual data (top), and representative mouse trajectories during the probe phase. **J**, Schematic of the TCT paradigm. **K**, *Left*: Time spent in the chamber containing the object (o) and the conspecific (m) during the second phase of the TCT. *Right*: Computed social discrimination index as shown in I. **L**, *Left*: Time spent in the chamber containing the familiar (f) and novel mouse (n) during the third phase of the TCT. *Middle*: Computed social discrimination as shown in I and K. *Right*: Representative mouse trajectories. **M**, Schematic of the contextual FC paradigm. **N**, Percentage of time spent freezing in response to shocks during the fear learning phase of the FC. **O**, Percentage of time spent freezing during the fear retrieval, shown as cumulative fraction as in I, K, and L. **P**, Schematic of the auditory startle and PPI paradigms. **Q**, Startle responses to auditory pulses of increasing intensity. **R**, PPI of the auditory startle response for 4 pre-pulse intensities. **S**, Schematic of the USV paradigm. **T,** Number of total calls (left), fraction of jump calls (middle) and number of call types (right) in 10 min USV recordings from pairs of control and A-α7^KO^ mice. **U**, representative 10 min USV recordings. **V**, Integrated z-scores across all behaviors tested in eN-α7^KO^, iN-α7^KO^, and A-α7^KO^ mice relative to their controls. EPM: elevated plus maze. SP: sucrose preference. OF: open field. Unpaired Student’s *t*-test (C, D, I, K right, L right, O, T, U, V), Mann-Whitney test (E, G, L, V), paired Student’s *t*-test (K left, L left), Wilcoxon test (H, K, L left), two-way repeated measures ANOVA with Sidak’s multiple comparisons tests (N,R) and mixed-effects analysis (Q) were used.

A-α7^KO^ mice weighed normally (Fig.S7A), appeared healthy, and were indistinguishable from their same-sex control littermates when observed in their home-cage and upon handling. These mice performed normally in the Y-maze, showing rates of spontaneous alternations and number of arm visits identical to controls (Fig.3E and Fig.S6B), indicative of intact working memory, locomotor activity and general exploratory drive (also see below). Strikingly, however, A-α7^KO^ mice showed profound disruptions in all other MCCB-related assays. In the NOR (Fig.3F), A-α7^KO^ mice not only displayed lower discrimination index on the probe day (Fig.3I), but also showed a complete lack of preference towards the novel object (Fig.3H) despite having normal combined object exploration, locomotion, speed of ambulation, and exploratory behavior during the acquisition and probe trials (Fig.3G and Fig.S6C). This deficit could not be explained by novel object place preference either (Fig.S6C), together suggesting that this was a direct result of impaired attention, object feature discrimination, encoding and/or retrieval of this declarative or episodic-like form of memory^30^. Likewise, A-α7^KO^ mice had a reduced social drive, seen as a decreased preference for a conspecific over an inanimate object in the TCT (Fig.3J-K), and a complete lack of preference for the novel mouse during the social recognition phase (Fig.3J,L and Fig.S6D,E). This indicated impaired social recognition, social memory, and/or preference for social novelty. Since KO animals seemed capable of discriminating between the novel and familiar mouse during the initial 2 min of the social recognition assay (Fig.S6E), we interpret these results as a lack of preference towards social novelty rather than impaired social recognition/memory, overall consistent with a diminished social drive. In the FC assay, A-α7^KO^ mice showed an initial reduction in fear response to the shocks, echoing evidence implicating astrocytic α7nAChRs in pain^50^, but reached freezing levels identical to controls by the end of the shock protocol (Fig.3M,N). Nonetheless, they spent significantly less time freezing when reintroduced in the shock context 24 hours later (Fig.3N,O), but not when introduced in a neutral context (Fig.S6F), which is canonically interpreted as a deficit in contextual (spatial) fear learning and memory.

Although these tasks presumably probe distinct behavioral functions, the NOR, TCT, and FC rely on a two-phase process consisting, first, of an encoding phase, followed by a second exposure during which encoding is indirectly probed. This suggests that, aside from their reduced social drive, the phenotype of A-α7^KO^ mice could amount to a general learning and memory deficit. This prompted us to examine potential alterations in sleep architecture in A-α7^KO^ mice that could undermine overall memory consolidation, especially because astrocytes regulate slow wave sleep and sleep drive^51^, while cholinergic neuromodulation governs vigilance states^31^. However, we found no differences in the number or duration of wake, quiet-wake, non-REM, and REM bouts in the light (sleep) or the dark (active) phase, and no differences in the spectral content of any vigilance states in EEM/EMG studies (Fig.S7F-H), ruling out sleep disturbances as a common denominator for the observed behavioral deficits. Additionally, A-α7^KO^ mice exhibited noticeable changes in sensory gating and courtship ultrasonic vocalizations (USVs), two innate behaviors^27,52^. Indeed, KO animals showed an exaggerated PPI (Fig.3P,R), besides a normal startle response profile (Fig.3Q). Recordings of USVs produced by pairs of A-α7^KO^ mice evidenced a drastic reduction in the total number of calls and call complexity compared to controls, with significantly fewer jump calls that spanned distinct frequency bands and a shift towards simpler one-call bouts of USVs (Fig.3S-U). Altogether, these results outline general deficits caused by the loss of astrocytic α7nAChR in adulthood that were not observed in eN-α7^KO^ and iN-α7^KO^ mice, including strong spatial, semantic, and social cognition impairments that could be interpreted as broad learning/memory deficits, as well as reduced social drive and communication, and alterations in sensory gating (Fig.3V).

We next sought to test whether these deficits were specific to MCCB domains or reflected impairments across a broader behavioral spectrum. We thus probed A-α7^KO^ mice for signs of altered exploration/locomotion (OF, Fig.S7E), anxiety (EPM, Fig.S7B), pleasure-seeking (SPT, Fig.S7D), and nesting behavior (Fig.S7C) as we did for eN-α7^KO^ and iN-α7^KO^ mice. Shockingly, we found no differences between A-α7^KO^ mice and their controls in any of these assays, strongly suggesting that the deficits observed in A-α7^KO^ mice are specific to MCCB domains and ruling out anxiety, anhedonia, or gross motor/exploratory as causal factors (Fig.3V). Additionally, we found no genotype × sex interactions (Table1) indicating that the phenotype of A-α7^KO^ was not specifically driven by males or females besides occasional manifestations of sex-dependent strategies, which we interpret as the result of inherent sex-bias of mouse behavior paradigms^53^.

## Astrocytic α7nAChR contribution to behavior is D-serine dependent

Having demonstrated the central role of astrocytic α7nAChR in MCCB-relevant behaviors, we next sought to provide a mechanistic context to these observations. Cholinergic inputs govern several astrocyte activities that are consequential to circuit function, including the control of extracellular D-serine^38,40,50^, an astrocyte-derived obligatory agonist of synaptic NMDARs^54,55^. Owing to the importance of NMDARs to a myriad of cellular and molecular processes that form the basis of behavior and cognition, especially learning and memory, we reasoned that a deficit in D-serine availability (and, consequently, NMDAR function) could underlie some of the behavioral alterations observed in A-α7^KO^ mice. Adult A-α7^KO^ and control littermates were implanted with micro-dialysis probes in the hippocampus, a key structure for learning and memory, to allow for the continuous collection of interstitial fluid for 6 hours as mice freely behaved in an open field (Fig.4A). Analysis of the collected dialysates using high pressure liquid chromatography (HPLC), revealed that hippocampal D-serine levels were reduced by 50% in A-α7^KO^ mice compared to controls (Fig.4A), consistent with prior work linking astrocytic α7nAChR to D-serine availability^40,50,56^. To test whether this might be sufficient to hinder NMDAR-dependent functions, we examined NMDAR-dependent synaptic plasticity in acute hippocampal slices from adult A-α7^KO^ and control animals. We found that long-term synaptic potentiation (LTP) was diminished by 50% in A-α7^KO^ slices (Fig.4B, D) and could be restored to control-levels by exogenous D-serine (20 µM, Fig.4C, D). Together, these results suggest that A-α7^KO^ behavioral deficits might be caused by a D-serine deficiency. To test this directly, we supplemented the drinking water of A-α7^KO^ mice and littermate controls with D-serine for 7 days (1.5-2.4 mg/ml, equivalent to 250-400 mg/kg/day)^57^ prior to probing their performance at the NOR, TCT and FC. Remarkably, under these conditions, the deficits of A-α7^KO^ were fully restored. The performance of D-serine-treated A-α7^KO^ mice was indeed identical to that of D-serine-treated controls in all aspects (Fig.4E-H, S8), including social drive, strongly indicating that D-serine is a causal mediator in the behavioral effect of astrocytic α7nAChRs.

**Figure 4:**
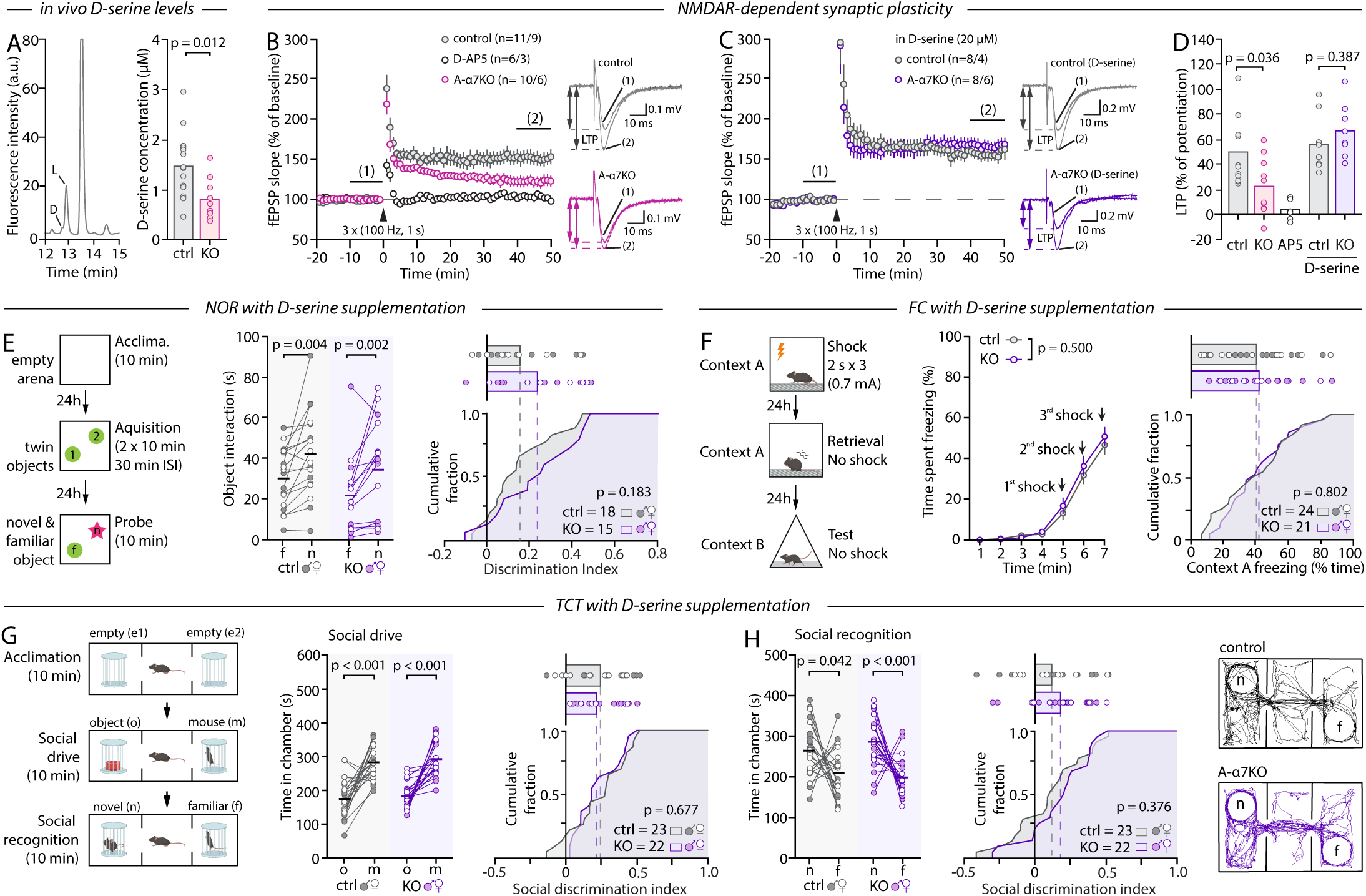
The phenotype of A-α7^KO^ mice is D-serine dependent. **A**, *Left*: Representative HPLC chromatograms. *Right*: D-serine levels in micro-dialysates collected from control and A-α7^KO^ mice. **B,** Time course of LTP in control and A-α7^KO^ slices in response to a high frequency stimulation (HFS) protocol and representative traces taken from the last 10 min of baseline (1) and last 10 min of post LTP induction (2). N-values indicdate the number of slice replicates and mouse replicates. **C**, Same as **B** in the presence of 20 µM D-serine applied 15 min prior to the HFS protocol. (**D**) Summary bar graphs showing the magnitude of LTP. **E**, *Left*: Schematic of the NOR paradigm. *Middle*: Time spent interacting with the novel and familiar object during the probe phase. *Right*: Computed object discrimination index of control and A-α7^KO^ mice supplemented with D-serine in their drinking water. **F**, *Left*: Schematic of the FC paradigm. *Middle*: Percentage of time spent freezing in response to shocks during the fear learning phase. *Right*: Percentage of time spent freezing during fear retrieval of the FC by control and A-α7^KO^ mice supplemented with D-serine. **G**, *Left*: Schematic of the TCT paradigm. *Middle*: Time spent in the chamber containing the object (o) and the conspecific (m) during the second phase of the TCT. *Right*: Computed social discrimination index (right) of control and A-α7^KO^ mice supplemented with D-serine. **H**, *Left*: Time spent in the chamber containing the familiar (f) and novel mouse (n) during the third phase of the TCT. *Right*: Computed social discrimination of control and A-α7^KO^ mice supplemented with D-serine, and representative trajectories. Unpaired Student’s *t*-test (A, D, E right, F right, H right), Mann-Whitney test (D, G right), paired Student’s *t*-test (E left, G left, H left), Wilcoxon test (E left), two-way repeated measures ANOVA with Sidak’s multiple comparisons tests (F middle) were used.

To further substantiate this finding, we took advantage of natural fluctuations in D-serine availability that occur throughout the sleep-wake cycle^40^. Prior work indeed showed that the astrocyte α7nAChR/D-serine pathway temporally integrates cholinergic tone: astrocyte α7nAChR-dependent D-serine secretion is high during the subjective day of mice (dark phase) and this signaling pathway is mostly inactive during the subjective night (light phase) when cholinergic activity is low^40^. Hence, we next tested whether the behavioral deficits of A-α7^KO^ would be masked if mouse behavior was probed in the light phase. That is, even though mouse behavior performance inherently varies throughout the day^58^, we reasoned that differences between controls and A-α7^KO^ mice would be concealed in the light phase when the astrocyte α7nAChR-D-serine signaling is naturally at its lowest in control mice^40^ and ablated in KOs. Consistent with this prediction, the performance of A-α7^KO^ mice and littermate controls at the NOR, TCT, and FC were indistinguishable when evaluated in the middle of the subjective sleep phase (ZT6, Fig.S9 and Fig.S10), confirming that astrocyte α7nAChRs support MCCB-related functions in a D-serine dependent manner. Importantly, this finding provides insight into why prior behavioral studies that examined constitutive astrocyte-specific α7nAChR KO mice in the light-phase found no deficits^20^ and could explain general inconsistencies in reports of behavioral phenotypes of constitutive α7nAChR KO mice^59^.

## The pro-cognitive effect of α7nAChR-therapeutics requires astrocytes

A multitude of α7nAChR agents have been tested in clinical studies in the past two decades to restore cognitive functions in schizophrenia, AD, and PD^8,9^. In contrast to their mode of action at the molecular level, which is often well-known, the mechanisms underlying the pro-cognitive properties of these drugs remain elusive. Having demonstrated that astrocytic rather than neuronal α7nAChRs are critical for behaviors assessed by the MCCB, we next tested whether astrocytes are a required substrate for the pro-cognitive potential of α7nAChR-targeting agents. We focused on the α7nAChR partial agonist EVP-6124 developed by FORUM Pharmaceuticals (Encenicline), because of its well-characterized mode of action on α7nAChR^60^ and its specificity (K_i_ = 9.98 nM). It is also of interest owing to its demonstrated clinical efficacy in healthy subjects, patients with AD, and patients with schizophrenia across all cognitive scales in Phase-II double-blind, randomized, placebo-controlled trials^61–64^, and because it is the only α7nAChR modulator to date to have been subsequently tested in Phase-III trials for the treatment of cognitive deficits in schizophrenia and AD (NCT01969123, NCT01969136, NCT01716975, NCT01714713, NCT01714661). Pharmacokinetic study of EVP-6124 in wild-type mice confirmed its rapid brain bioavailability, peaking at 30 min post-injection in males and 60 min in females, and its rapid clearance/metabolism, being undetectable in all animals within 24 hours (Fig.5A). We next tested its behavioral efficacy over a range of concentrations in wild-type mice completing the NOR task (Fig.5B). Consistent with clinical data^63,65^ and with its action as a partial agonist over a narrow range of concentrations^60^, EVP-6124 had a sharp dose-dependent effect on behavior performance in the light phase (ZT6, Fig.5C). Mice that received EVP-6124 at 0.3 mg/kg intraperitoneally (i.p.) prior to the acquisition phase of the NOR showed a 2-fold greater performance than vehicle-treated animals on the probe trial (Fig.5D). Importantly, this improvement was not the result of increased activity, increased time spent interacting with objects, or changes in object preference (Fig.S11), hence recapitulating the pro-cognitive effect of EVP-6124 from pre-clinical and clinical studies^60,63,65^. We next tested EVP-6124 in A-α7^KO^ and control mice at ZT6, when their performance at the NOR was identical (Fig.S9). Like in wild-type mice, EVP-6124 augmented behavior in control animals nearly 3-fold compared to vehicle-treatment, with no changes in activity or exploration (Fig.5E and Fig.S11). However, EVP-6124 was ineffective when administered to A-α7^KO^ mice: vehicle- and EVP-6124-treated A-α7^KO^ mice showed identical object discrimination (Fig.5F,G). Given these results, we conclude that astrocytic α7nAChRs are required for the pro-cognitive effect of a flagship α7nAChR drug tested in Phase-III clinical trials in recent years.

**Figure 5:**
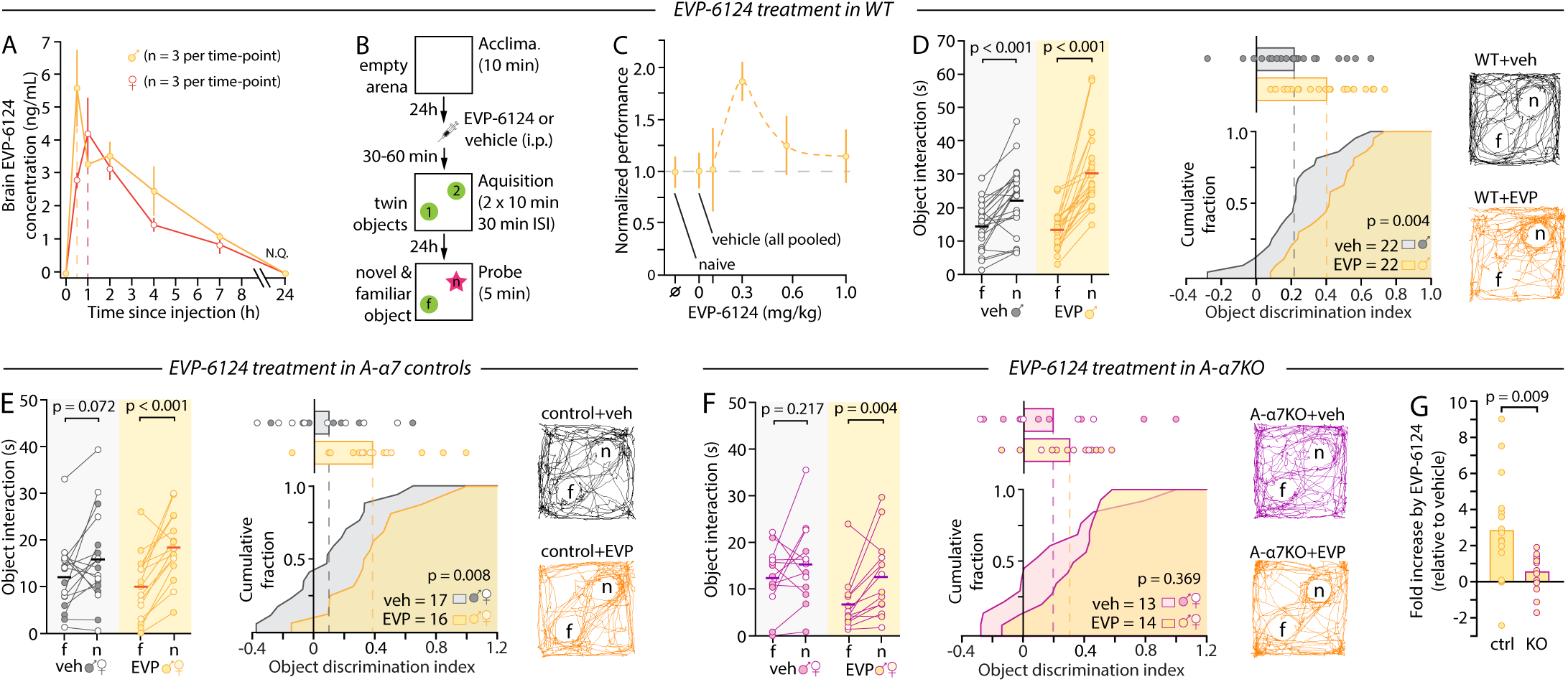
The pro-cognitive enhancer EVP-6124 requires astrocytic α7nAChR to augment behavior. **A**, Pharmacokinetic studies of EVP-6124 in the brain of wild-type male and female mice after i.p. administration (0.3 mg/kg in 0.36% DMSO in saline, i.p. at time 0). N.Q.: non quantifiable. **B**, Schematic of the NOR paradigm and EVP-6124 injection timeline. **C**, Dose-response curve of EVP-6124 (0.1, 0.3, 0.6 and 1.0 mg/kg) on the performance in the NOR relative to vehicle-treated cage-mates. Naïve: untreated animals. Vehicle: all vehicle-treated animals (n=97). The performance of EVP-6124-treated animals was normalized to that of their respective vehicle-treated cage mates (n = 16 to 34 per group). **D,** *Left*: Time spent interacting with the novel (n) and familiar (f) object during the probe phase of the NOR by mice treated with vehicle or 0.3mg/kg EVP-6124. *Right*: Computed object discrimination index and representative trajectories. **E, F**, same as D in control (E) and A-α7^KO^ (F) mice. **G**, Bar graphs summarizing the enhancing effect of EVP-6124, relative to vehicle treatment, in control and A-α7^KO^ mice. Unpaired Student’s *t*-test (D right, E right, F right, G), paired Student’s *t*-test (D left, E left, F left), Wilcoxon test (D left, F left), Kruskal-Wallis test with Dunn’s test (C) were used.

## Discussion

Although the functional enhancement of α7nAChRs with therapeutic agents has solidified the pro-cognitive value of the receptor in rodents^7,8,10^ and humans^7,63^, the substrate and mechanistic by which α7nAChRs support cognition have remained elusive. By systematically assessing behavior phenotypes in three cell-type-specific inducible α7nAChR KO lines across 12 behavioral paradigms and 37 metrics, our findings demonstrate a central role of astrocytic rather than neuronal α7nAChRs in cognitive behaviors typically evaluated in schizophrenia trials. The loss of α7nAChRs in principal neurons markedly reduced whole-brain *Chrna7* expression but was overall inconsequential to behavior, and the loss of α7nAChRs in inhibitory neurons had minimal impact on MCCB-related cognitive behaviors, mainly causing alterations in other domains. In contrast, ablating α7nAChRs from astrocytes strongly impacted mouse performance in six out of seven MCCB-related tasks but left other behaviors completely unaffected. An important consideration in these experiments is that all three KO lines featured incomplete penetrance of recombination (∼50-70%), raising the possibility that the behavioral importance of astrocytic α7nAChRs is underestimated in our study. It also raises the possibility that the general lack of phenotype in eN-α7^KO^ and iN-α7^KO^ animals reflects the deletion of α7nAChR from an insufficient contingent of neurons. We consider this unlikely, however, in view of the clear evidence that *Chrna7* was successfully deleted in 50% or more of the desired cell-types. In addition, and remarkably, the behavioral alterations observed in all three lines, when compounded, largely replicate the constellation of phenotypes described in constitutive α7nAChR KOs over the years^32,66–68^. Together, these results echo a growing body of literature describing astrocytes as key computational units and underscore the need to systematically consider astrocyte signaling when dissecting the cellular and molecular correlates of behavior^69^.

Our findings link the pro-cognitive value of α7nAChRs to D-serine/NMDAR signaling: A-α7^KO^ behavioral alterations coincide with diminished D-serine levels in vivo, can be rescued by D-serine supplementation, and mirror daily changes in D-serine availability. In addition, the behavior phenotype of A-α7^KO^ mice remarkably mimics that of mice lacking the D-serine synthesis enzyme serine racemase^70,71^, and that of mice with NMDAR hypofunction^73,74^ or hypomorphic for the GluN1 subunit of NMDARs on which D-serine binds^75^. Collectively, this evidence strongly points to the astrocytic α7nAChR/D-serine/NMDAR pathway as a central mechanism underlying the pro-cognitive nature of α7nAChR signaling. In fact, reports consistently indicate that D-serine levels are decreased in the plasma or CSF of patients with schizophrenia^76–78^. In addition, supplementations with D-serine or its functional analogues (glycine or D-cyclo-serine) received considerable attention in past clinical attempts to improve cognition in schizophrenia^79^ (see NCT00322023, NCT00455702, NCT00000372). Although D-serine was not measured in trials of α7nAChR-targeting pro-cognitive drugs, our findings suggest that these parallel efforts converge on a shared α7nAChR/D-serine/NMDAR pathway.

The discovery that a Phase-III α7nAChR partial agonist requires astrocytic α7nAChRs to augment behavior in mice adds to recent human evidence linking the cognitive deficits of schizophrenia to α7nAChRs and glia^80^ and has important translational implications. Indeed, α7nAChR partial agonists like EVP-6124 are therapeutically constrained by their sharp dose-limiting efficacy and gastrointestinal side-effects^9^. D-serine and its functional analogues, on the other hand, are limited by their nephrotoxicity and poor brain bioavailability^81^. Identifying intracellular processes linking α7nAChR signaling to D-serine secretion in astrocytes might thus provide more exploitable and targeted therapeutic avenues to ameliorate cognitive aspects of psychiatric disorders. At this time, this is hindered by our fragmented knowledge of the rules and mechanisms of input-output transformation in astrocytes, emphasizing that a better understanding of these cells has become critical for translational research.

A potential limitation to acknowledge in this study is that Cre was expressed in a subset of newborn neurons of the DG in A-α7^KO^ mice, which could have contributed to their behavioral deficits. The reason this is unlikely is 6-fold. First, only 12% of DG neurons were Cre-expressing. Second, adult newborn neurons in the DG are GABAergic and express Gad2^82^, yet the phenotype of iN-α7^KO^ and A-α7^KO^ are clearly different. Third, ∼50% of these DG newborn neurons are eliminated within 4 weeks^83^. Fourth, the functional integration of the surviving neurons into the local circuitry is complete 7-8 weeks after cell birth, which is beyond our behavioral timeline. Fifth, the phenotype of A-α7^KO^ mice was time-of-day dependent and, sixth, rescued by D-serine, both of which are consistent with an astrocyte-driven effect, but inconsistent with a newborn-neuron-mediated effect. In total, we thus genuinely interpret the behavioral phenotype of A-α7KO mice as driven by the ablation of α7nAChR from astrocytes.

In summary, our findings challenge the common view that neurons are the cellular substrate of cognition and sole target of pro-cognitive therapeutics, and further underscore how insights from astrocyte biology could inform new treatments for psychiatric disorders.

## Supplementary figure legends

**Figure S1:**
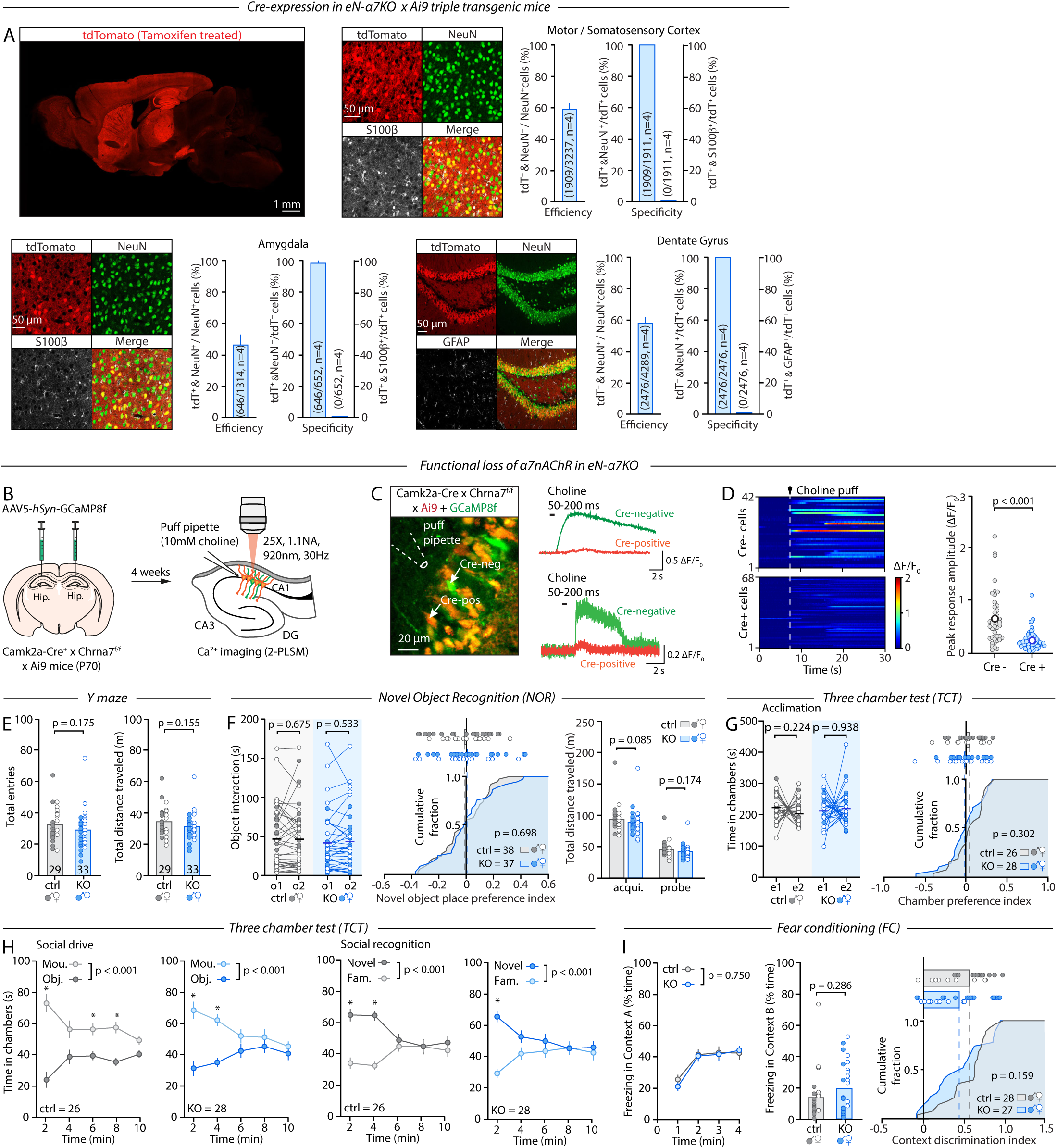
eN-α7^KO^ validation and cognitive behavior. **A**, IHC quantification of specificity and efficiency of recombination in eN-α7^KO^ ⁢ Ai9 mice in the motor cortex and somatosensory cortex (pooled), amygdala, and dentate gyrus. Shown are quantifications of efficiency and specificity with the pan-neuronal marker NeuN. **B**, schematic of the experimental paradigm to validate the loss of functional of α7nAChR in Cre-expressing neurons in the hippocamps of eN-α7^KO^ mice. GCaMP8f expressing neurons were considered Cre-positive is they also expressed tdTomato (indicative of Cre-induced recombination of the Ai9 allele). **C**, *Left*: Representative 2-PLSM field of view showing a majority of neurons expressing both tdTomato and GCaMP8f (yellow). A few neurons are unrecombined and solely express GCaMP8f (green). *Right*: Two sets of representative fluorescence time-series from Cre-positive and Cre-negative cells (shown in the left 2-PLSM image) in response to puff applications of choline. **D**, Kymograph (left) and individual peak or choline-evoked responses (right) of 42 Cre-negative and 68 Cre-positive neurons. **E**, Number of arm entries (left) and total distance traveled (right) in the Y-maze. **F**, Time spent exploring the two identical objects during the acquisition phase of the NOR (left), computed object place preference during the acquisition phase, based on position where the novel object is placed on the subsequent phase (middle), and total distance traveled during the acquisition and probe phases (right). **G**, Time spent in the two empty side chambers during the first phase of the TCT (left) and computed chamber preference index (right). **H**, Temporal profile of mouse vs object chamber exploration during the phase 2 (social preference) and novel vs. familiar mouse chamber during phase 3 (social memory) of the TCT. **I**, Time course of the percentage of time spent freezing during the probe phase of the FC (left), percentage of time spent freezing in context B (middle), and computed context discrimination index (right). Unpaired Student’s *t*-test (F middle, G right, I right), Mann-Whitney test (D, E left, E right, F right, I middle), paired Student’s *t*-test (G left), Wilcoxon test (F left, G left), two-way repeated measure ANOVA with Sidak’s multiple comparisons test (H, I left) were used.

**Figure S2:**
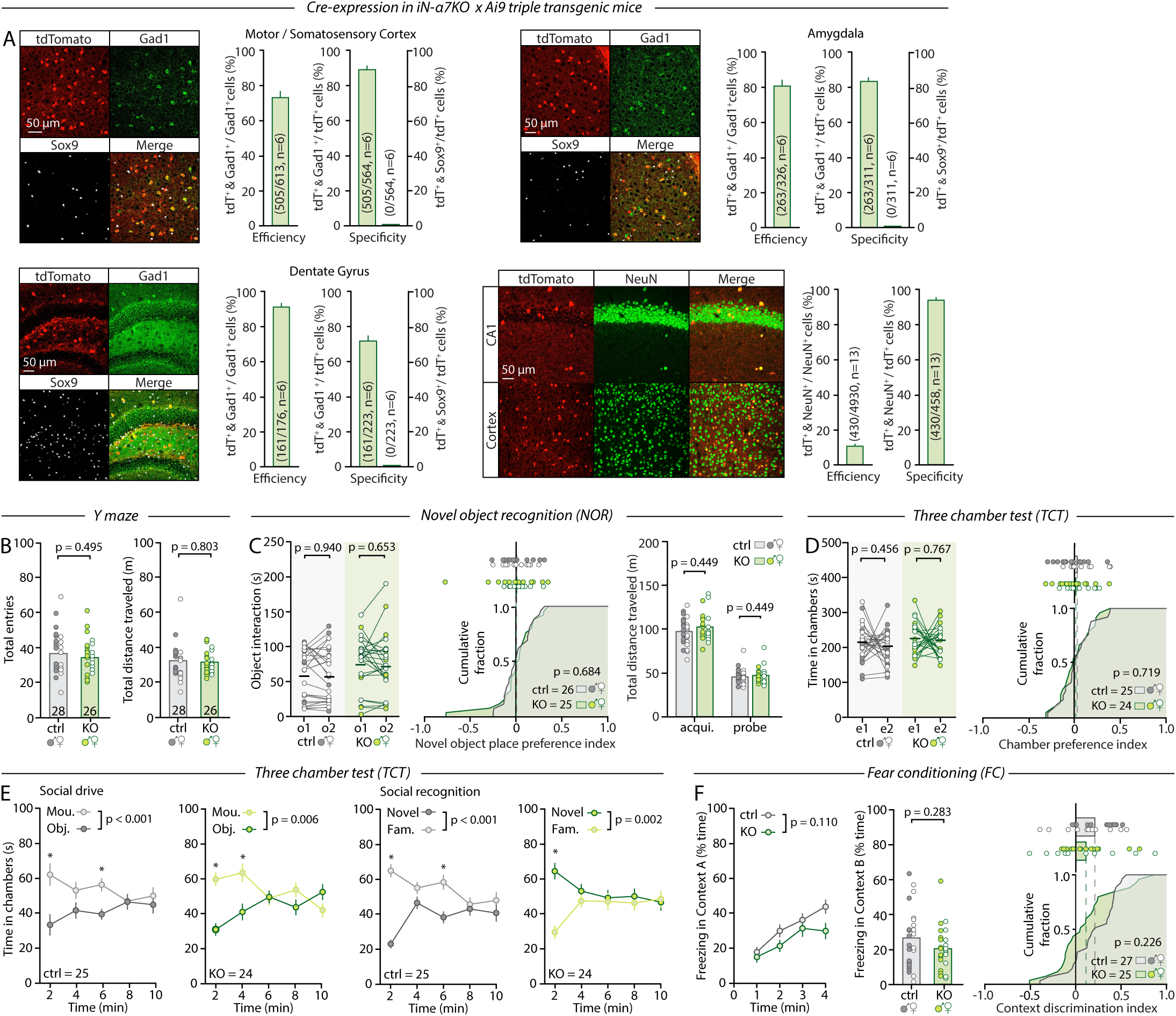
iN-α7^KO^ validation and cognitive behavior. **A**, IHC quantification of specificity and efficiency of recombination in iN-α7^KO^ ⁢ Ai9 mice in the somatosensory cortex and motor cortex (pooled), amygdala and dentate gyrus. Shown are quantifications of efficiency and specificity with the pan-neuronal marker NeuN, and with the GABAergic neuron marker Gad1. **B**, Number of arm entries (left) and total distance traveled (right) in the Y-maze. **C**, Time spent exploring the two identical objects during the acquisition phase of the NOR (left), computed object place preference during the acquisition phase, based on the position where the novel object is placed on the subsequent phase (middle), and total distance traveled during the acquisition and probe phases (right). **D**, Time spent in the two empty side chambers during the first phase of the TCT (left) and computed chamber preference index (right). **E**, Temporal profile of mouse vs object chamber exploration during phase 2 (social preference) and novel vs familiar mouse chamber during phase 3 (social memory) of the TCT. **F**, Time course of the percent of time spent freezing during the probe phase of the FC (left), percentage of time spent freezing in context B (middle), and computed context discrimination index (right). Unpaired Student’s *t*-test (B left, D right, F right), Mann-Whitney test (B right, C right, F middle), paired Student’s *t*-test (C left, D left), Wilcoxon test (C left), two-way repeated measures ANOVA with Sidak’s multiple comparisons test (E, F left) were used.

**Figure S3:**
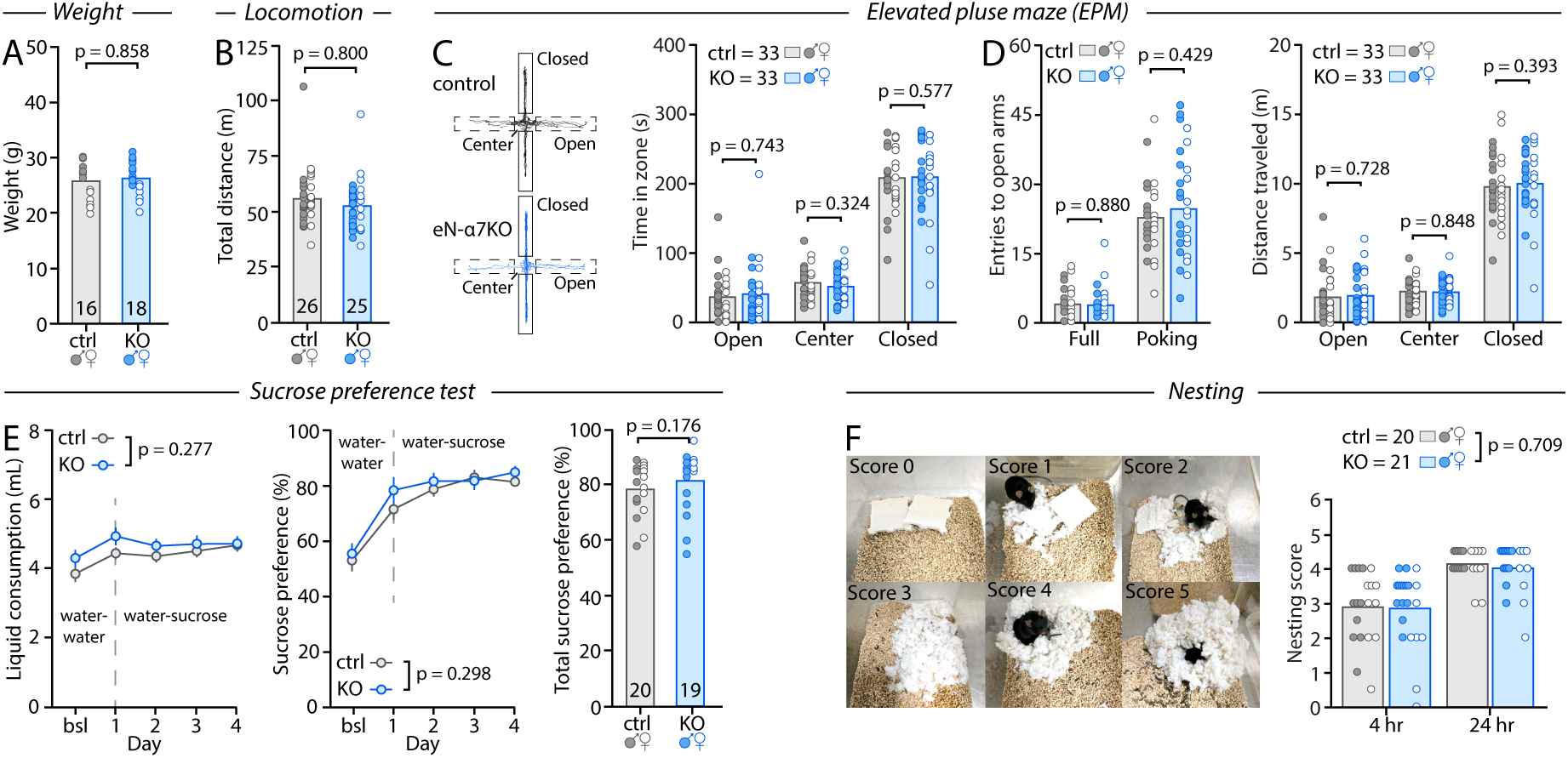
Non-cognitive behavior performance of eN-α7^KO^ mice. **A**, Body weight of control and eN-α7^KO^ mice at P90-100. **B**, Distance traveled in 10min in an empty open field under dim lighting (230 lux). **C**, EPM data showing representative trajectories (left) and the amount of time spent in the open arms, closed arms and center of the maze (right). **D**, *Left*: Number of entries in the open arms by type (full vs poking, left). *Right*: Total distance traveled in the open arms, closed arms and center of the maze. **E**, Sucrose preference test showing the combined fluid intake from both bottles per day (left), the sucrose preference calculated as the percentage of total daily fluid consumed from the sucrose bottle (middle), and mean sucrose preference over the final 3 days of testing (right). Bottle positions were alternated daily to control for side bias. Bsl: baseline (water–water) intake averaged over the 3 days prior to sucrose presentation. **F**, *Left*, representative illustrations of nest completion and their associated scores. *Right*, Nesting scores 4-6 hours and 24 hours after mice were given 5g of intact bedding material. Unpaired Student’s *t*-test (D left), Mann-Whitney test (A, B, C, D left, D right, E right), two-way repeated measure ANOVA with Sidak’s multiple comparisons test (E left, E middle, F) were used.

**Figure S4:**
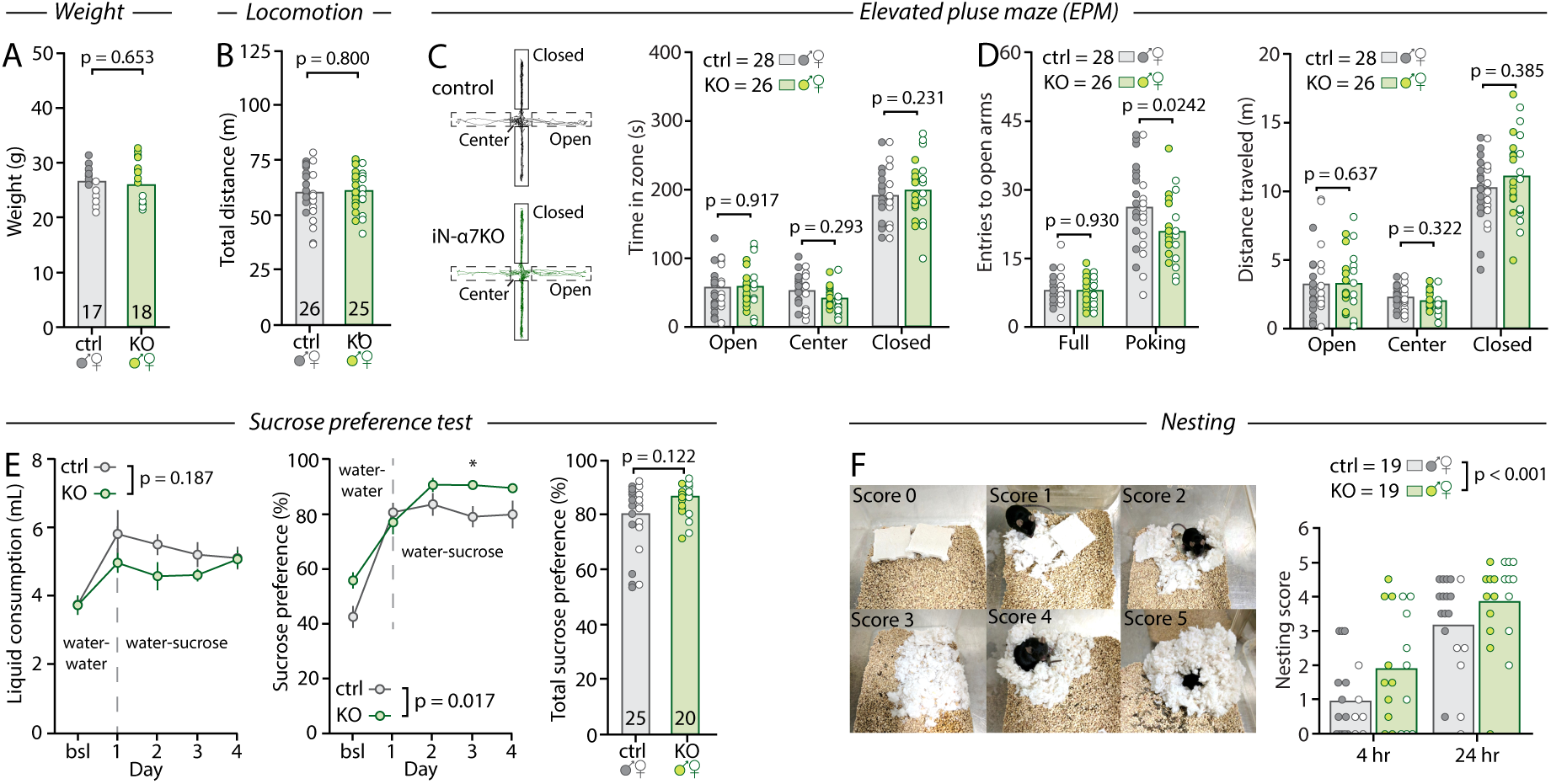
Non-cognitive behavior performance of iN-α7^KO^ mice. **A**, Body weight of control and iN-α7^KO^ mice at P90-100. **B**, Distance traveled in 10 min in an empty open field under dim lighting (230 lux). **C**, EPM data showing representative trajectories (left) and the amount of time spent in the open arms, closed arms and center of the maze (right). **D**, *Left*: Number of entries in the open arms by type (full vs poking, left). *Right*: Total distance traveled in the open arms, closed arms and center of the maze. **E**, Sucrose preference test showing the combined fluid intake from both bottles per day (left), the sucrose preference calculated as the percentage of total daily fluid consumed from the sucrose bottle (middle), and mean sucrose preference over the final 3 days of testing (right). Bottle positions were alternated daily to control for side bias. Bsl: baseline (water–water) intake averaged over the 3 days prior to sucrose presentation. **F**, *Left*, representative illustrations of nest completion and their associated scores. *Right*, Nesting scores 4-6 hours and 24 hours after mice were given 5 g of intact bedding material. Unpaired Student’s *t*-test (B, C, D left), Mann-Whitney test (A, D right, E right), two-way repeated measures ANOVA with Sidak’s multiple comparisons test (D left, F) were used.

**Figure S5:**
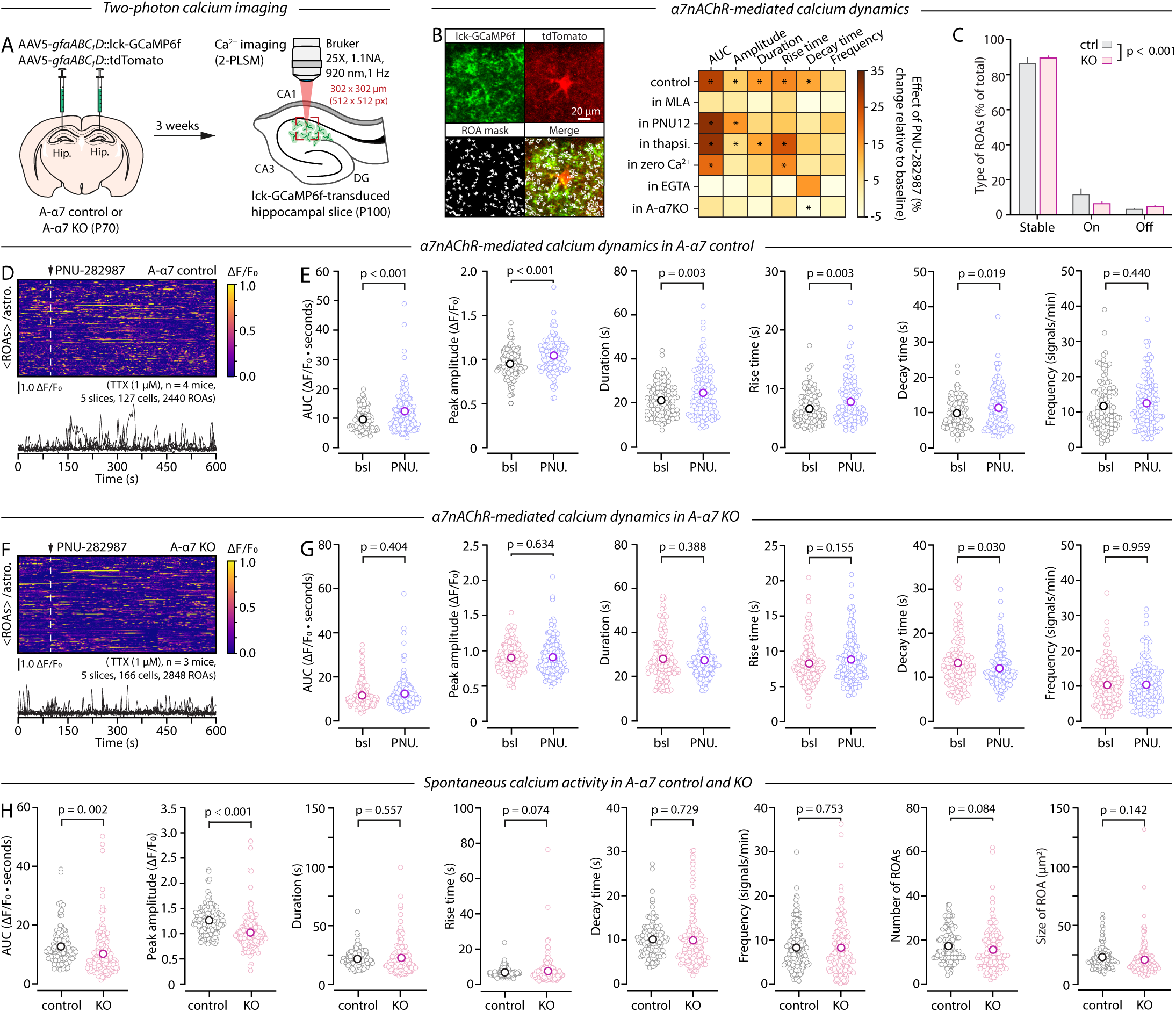
α7nAChRs on astrocytes control Ca^2+^ dynamics. **A**, Schematic of experimental design for the expression lck-GCaMP6f in astrocytes and subsequent 2-PLSM recordings of Ca^2+^ activity in brain slices. **B**, *Left:* Representative fluorescence images of lck-GCaMP6f in a tdTomato-expressing astrocyte and map of regions of activity (ROAs) captured over a 10 min recording. *Right:* Summary of the effect of PNU-282987, a specific α7nAChR agonist, on the properties of Ca^2+^ transients across conditions. Results are shown as average percentage change relative to baseline. **C**, Proportions of ROA types. Stable: active ROAs prior and during PNU-282987 application. On: ROAs exclusively active during the PNU-282987 epoch. Off: ROAs that were active during baseline but inactive upon PNU-282987 application. **D**, Kymograph showing the effect of PNU-282987 (500 nM) on astrocyte Ca^2+^ activity (shown as cell average of ROAs) in hippocampal slices from control animals and representative ΔF/F0 traces. Vertical dotted line shows time of bath application of PNU-282987. **E**, Pair-wise data showing the effect of PNU-282987 on the area under the curve (AUC), peak amplitude, kinetics and frequency of Ca^2+^ transients in astrocytes from control mice. **F, G**, Same as E and D in A-α7^KO^ mice. **H**, Comparison of the AUC, peak amplitude, kinetics and frequency of spontaneous Ca^2+^ transients in astrocytes of control and A-α7^KO^ mice, as well as number and size of ROAs per cell. Permutation test (B, E, G, H) and Chi-squared test (C) were used.

**Figure S6:**
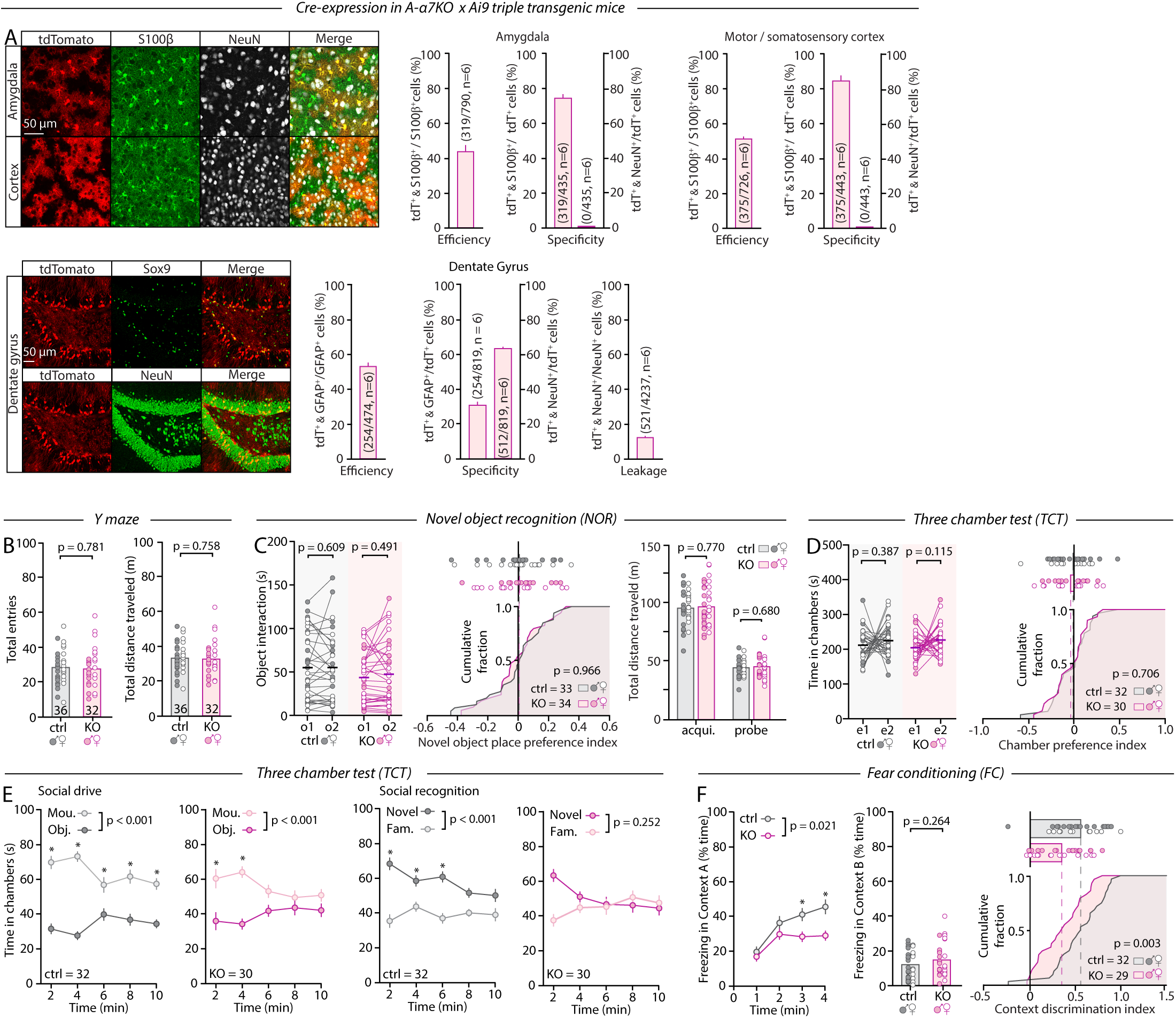
A-α7^KO^ validation and cognitive behavior. **A**, IHC quantification of specificity and efficiency of recombination in A-α7^KO^ ⁢ Ai9 mice in the motor cortex and somatosensory cortex (pooled), amygdala and dentate gyrus. Shown are quantifications of efficiency and specificity with the astrocyte markers GFAP, S100β or Sox9 and the neuronal marker NeuN. Note the lack of specificity in the DG where only ∼60% of the Cre-expressing cells are neurons (∼35% astrocytes), corresponding to a leakage of Cre-expressing in ∼12% of all neurons (see main text) **B**, Number of arm entries (left) and total distance traveled (right) in the Y-maze. **C**, Time spent exploring the two identical objects during the acquisition phase of the NOR (left), computed object place preference during the acquisition phase, based on position where the novel object is placed on the subsequent phase (middle), and total distance traveled during the acquisition and probe phases. **D**, Time spent in the two empty side chambers during the first phase of the TCT (left) and computed chamber preference index (right). **E**, Temporal profile of mouse vs object chamber exploration during phase 2 (social preference) and novel vs familiar mouse chamber during phase 3 (social memory) of the TCT. **F**, Time course of the percentage of time spent freezing during the probe phase of the FC (left), percentage of time spent freezing in context B (middle), and computed context discrimination index (right). Unpaired Student’s *t*-test (B left, B right, C middle, C right, D right, F right), Mann-Whitney test (F middle), paired Student’s *t*-test (D left), Wilcoxon test (C left), two-way repeated measure ANOVA with Sidak’s multiple comparisons test (E, F left) were used.

**Figure S7:**
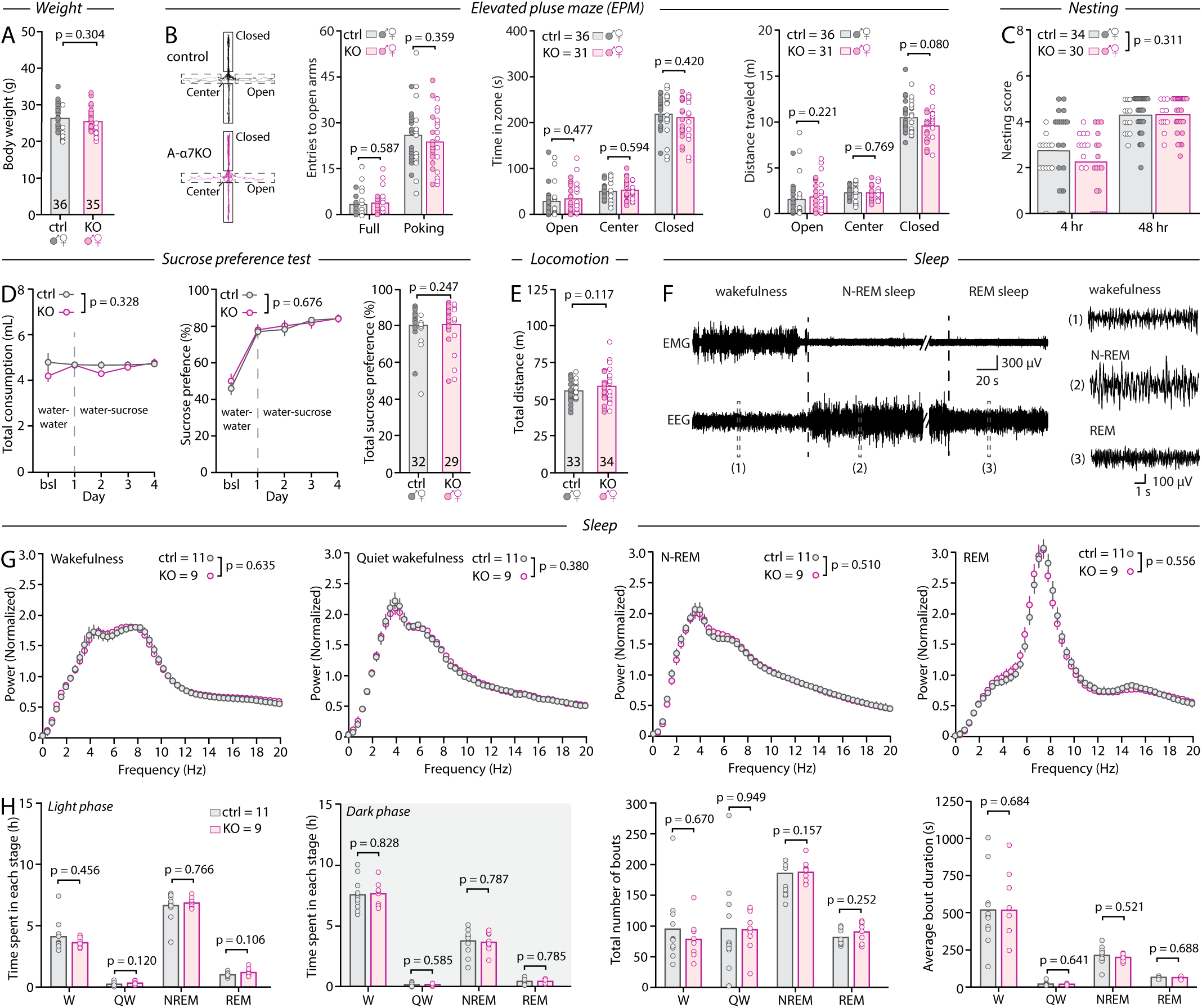
Performance of A-α7^KO^ mice in the non-cognitive domain. **A**, Body weight of control and A-α7^KO^ mice at P90. **B**, EPM data showing representative trajectories and the number of entries in the open arms by type (full vs. poking), the amount of time spent in the open arms, closed arms and center of the maze, and the total distance traveled in the open arms, closed arms and center of the maze. **C**, Nesting scores 4-6 hours and 24 hours after mice were given 5 g of intact bedding material. **D**, Sucrose preference test showing the combined fluid intake from both bottles per day (left), the sucrose preference calculated as the percentage of total daily fluid consumed from the sucrose bottle (middle), and mean sucrose preference over the final 3 days of testing (right). Bottle positions were alternated daily to control for side bias. Bsl: baseline (water–water) intake averaged over the 3 days prior to sucrose presentation. **E**, Distance traveled in 10 min in an empty open field under dim lighting (230 lux). **F**, Representative examples of time-locked EEG and EMG traces and expanded examples of wake, N-REM and REM bouts. G, Power spectra of Wake, quiet wake, N-REM and REM EEG traces. Spectra were normalized across the entire frequency band and averaged. **H**, *Left*: Time spent in each vigilance state in the light (sleep) and dark (active) phase. *Middle*: Total number of bouts spent in each vigilance state. *Right*: Average bouts duration. Unpaired Student’s *t*-test (A, B left, B middle, E, H), Mann-Whitney test (B right, D right, H), two-way repeated measure ANOVAs with Sidak’s multiple comparisons test (C, D left, D middle, G) were used.

**Figure S8:**
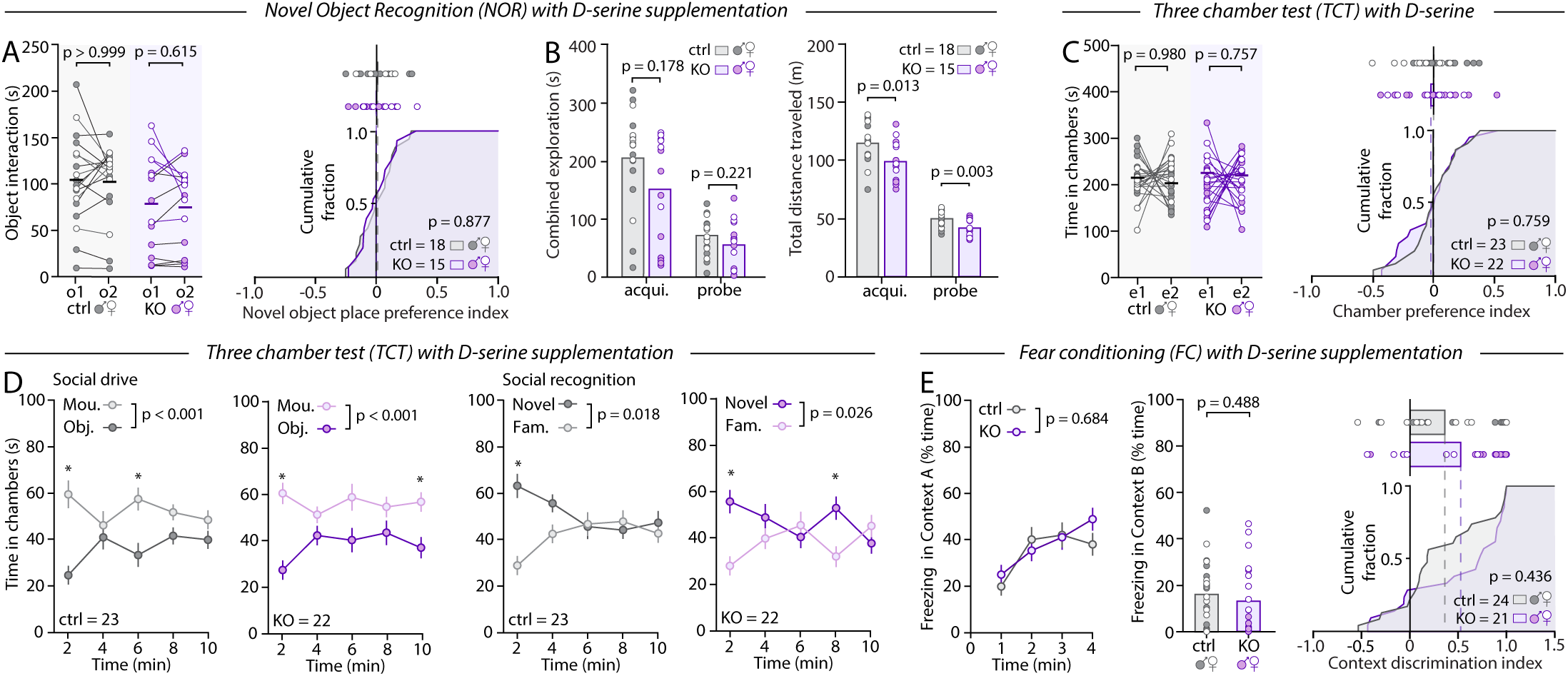
The phenotype of A-α7^KO^ mice is D-serine dependent. **A**, Time spent exploring the two identical objects during the acquisition phase of the NOR (left), computed object place preference during the acquisition phase based on the position where the novel object is placed during the subsequent phase (right). B, Total exploration time (left) and total distance traveled (right) during the acquisition and probe phases, by control and A-α7^KO^ mice supplemented with D-serine for 7 days. **C**, Time spent in the two empty side chambers during the first phase of the TCT (left) and computed chamber preference index (right) for control and A-α7^KO^ mice supplemented with D-serine for 7 days. **D**, Temporal profile of mouse vs object chamber exploration during phase 2 (social preference) and novel vs familiar mouse chamber during phase 3 (social memory) of the TCT by control and A-α7^KO^ mice supplemented with D-serine for 7 days. **E**, Time course of the percent of time spent freezing during the probe phase of the FC (left), percent of time spent freezing in context B (middle), and computed context discrimination index (right) for control and A-α7^KO^ mice supplemented with D-serine for 7 days. Unpaired Student’s *t*-test (A right, B left, B right, C right), Mann-Whitney test (B left, E middle, E right), paired Student’s *t*-test (A left, C left), two-way repeated measure ANOVA with Sidak’s multiple comparisons test (D, E left) were used.

**Figure S9:**
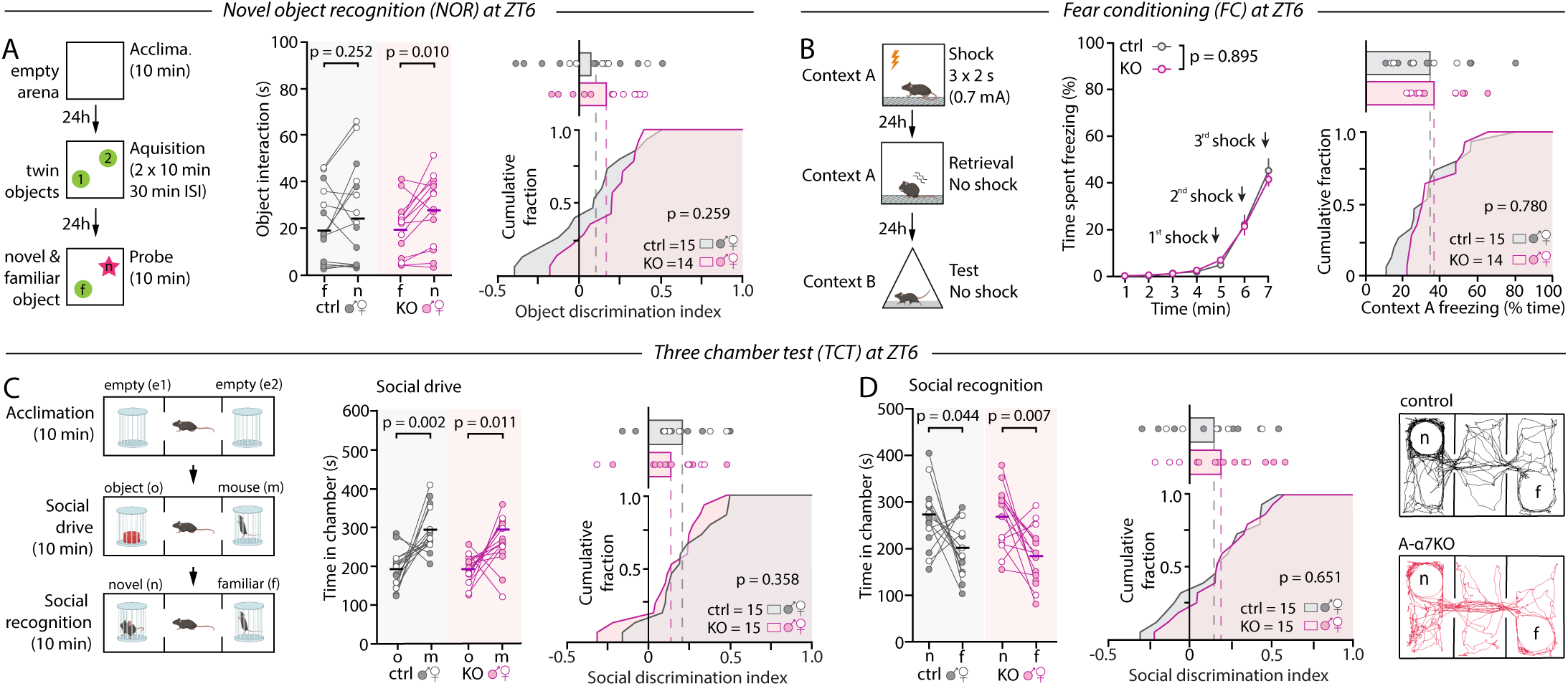
The phenotype of A-α7^KO^ mice is concealed at ZT6. **A**, Schematic of the NOR paradigm (left), time spent interacting with the novel and familiar object by control and A-α7^KO^ mice during the probe phase of the NOR at ZT6 (middle) and computed object discrimination index (right). **B**, Schematic of the FC paradigm (left), percentage of time spent freezing in response to shocks during the fear learning phase of the FC at ZT6 (middle), and percentage of time spent freezing during fear retrieval at ZT6 shown as a cumulative fraction. **C**, Schematic of the TCT paradigm (left), time spent in the chamber containing the object (o) and the conspecific (m) during the second phase of the TCT at ZT6 (middle) and computed social discrimination index (right). **D**, Time spent in the chamber containing the familiar (f) and novel mouse (n) during the third phase of the TCT at ZT6 (left), computed social discrimination (middle) and representative trajectories (right). Unpaired Student’s *t*-test (A right, C right, D right), Mann-Whitney test (B right), paired Student’s *t*-test (A left, C left, D left), two-way repeated measure ANOVAs with Sidak’s multiple comparisons test (B left) were used.

**Figure S10:**
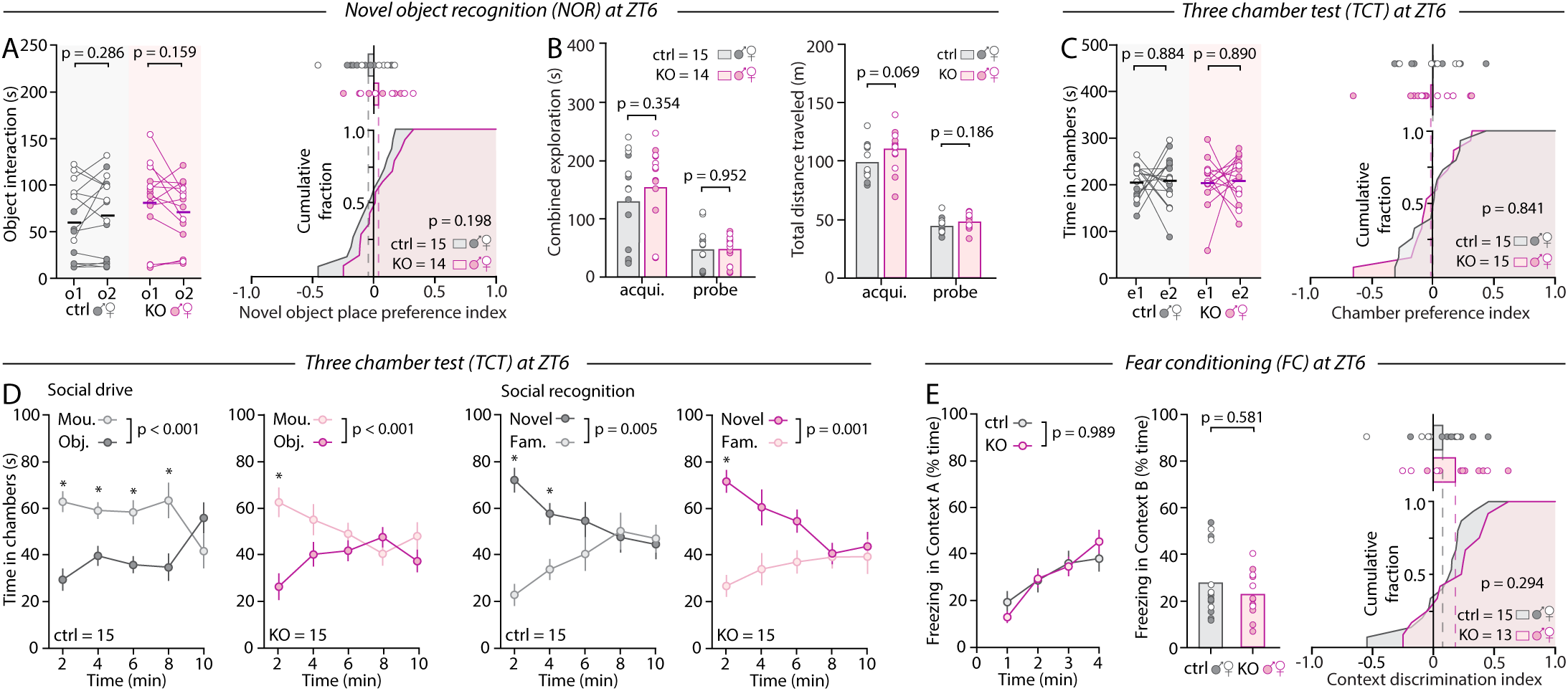
Control and A-α7^KO^ behavior at ZT6. **A**, Time spent exploring the two identical objects during the acquisition phase of the NOR at ZT6 (left), and computed object place preference during the acquisition phase based on position where the novel object is placed on the subsequent phase (right). **B**, Total object exploration (left) and total distance traveled (right) during the acquisition and probe phases at ZT6. **C**, Time spent in the two empty side chambers during the first phase of the TCT (left) and computed chamber preference index (right) at ZT6. **D**, Temporal profile of mouse vs object chamber exploration during the phase 2 (social preference) and novel vs familiar mouse chamber during the phase 3 (social memory) of the TCT at ZT6 by control and A-α7^KO^ mice. **E**, Time course of the percentage of time spent freezing during the probe phase of the FC (left), percentage of time spent freezing in context B (middle), and computed context discrimination index (right) for control and A-α7^KO^ at ZT6. Unpaired Student’s *t*-test (A right, B left, B right, C right, E right), Mann-Whitney test (B left, E middle), paired Student’s *t*-test (A left, C left), Wilcoxon test (C left), two-way repeated measure ANOVAs with Sidak’s multiple comparisons test (D, E left) were used.

**Figure S11:**
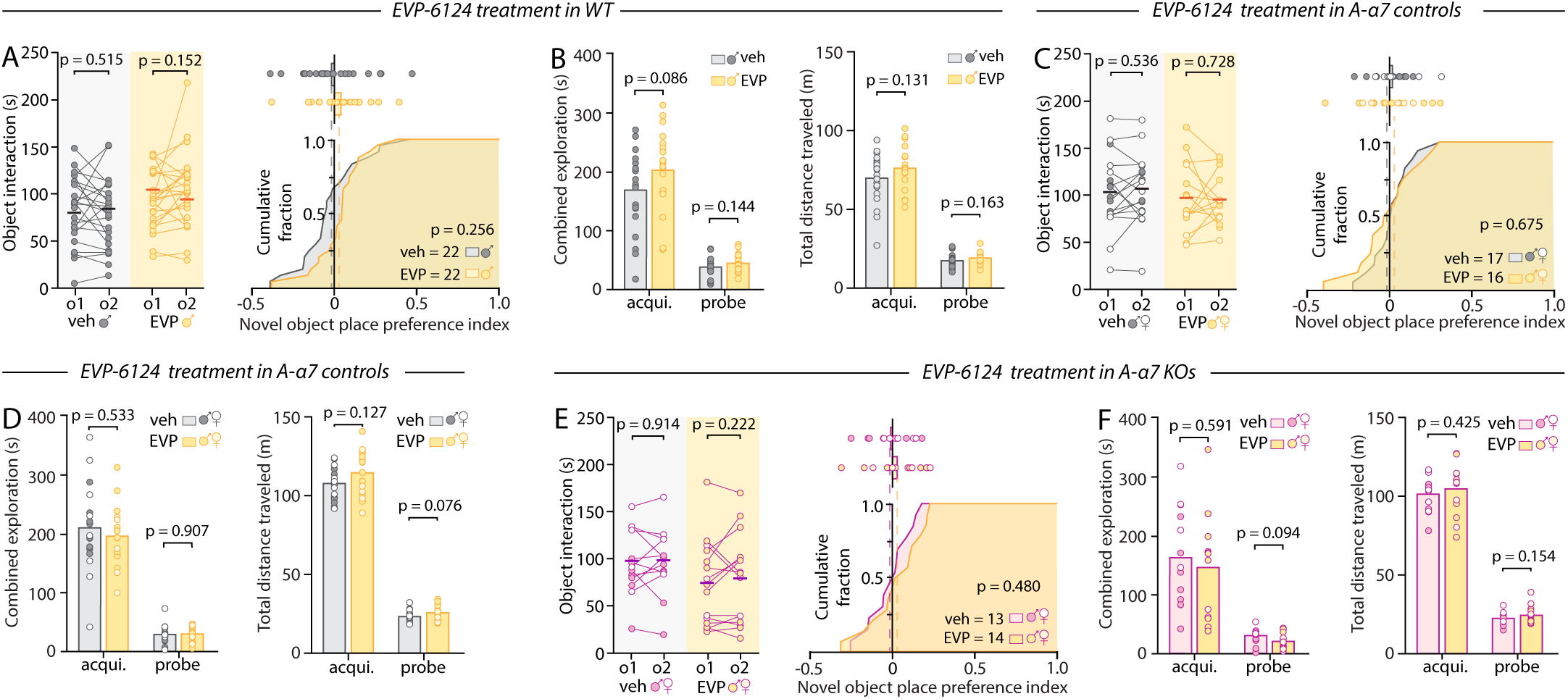
Effect of EVP-6124 in wild-type, control and A-α7^KO^ mice. **A**, Time spent exploring the two identical objects during the acquisition phase of the NOR (left) and computed object place preference during the acquisition phase based on the position where the novel object is placed on the subsequent phase (right) by wild-type mice administered EVP-6124 (0.3 mg/kg, i.p.) or vehicle (0.36% DMSO). **B**, Time spent exploring both objects combined during the acquisition and probe phase of the NOR (left), and total distance traveled during the acquisition and probe phase (right), by wild-type mice administered EVP-6124 or vehicle. **C, D**, Same as A and B for A-α7 control mice. **E, F**, Same as A and B for A-α7^KO^ mice. Unpaired Student’s *t*-test (A right, B left, B right, C right, D left, D right, E right, F left, F left), paired Student’s *t*-test (A left, C left, E left) were used.

## Acknowledgments

This work was supported by the National Institutes of Health (grant R01MH127163-01 to T.P.); the Department of Defense (grant W911NF-21-1-0312 to T.P.); the Brain & Behavior Research Foundation (NARSAD Young Investigator Award 28616 to T.P.), the Whitehall Foundation (grant 2020-08-35 to T.P.); the McDonnell Center for Cellular and Molecular Neurobiology (grant 22-3930-26275U to T.P.); and the Center for Drug Discovery at Washington University in St Louis (. The authors thank Dr. Peter Bayguinov (Washington University Center for Cellular Imaging), and Dr. Carla Yuede and Dr. Susan Maloney (Animal Behavior Core) for their critical expertise, and Dr. Gareth Rurak and Hallie Youker for their comments on the manuscript.

## Author contributions

Conceptualization: Y.W., M.T. T.P.; Funding acquisition: T.P.; Methodology: Y.W., M.T. Y.D., S.W., T.P.; Investigation: Y.W., M.T., Y.D., S.W., H.A., K.B.L., H.A., R.M.; Project administration: T.P.; Supervision: Y.W., P.G.H., T.P.; Visualization: Y.W., T.P.; Writing – original draft: Y.W.; Writing – review & editing: all authors.

## Data and materials availability

All raw data will be made available on a public repository by the time of publication. No codes were generated for this study. All codes and methods for the STARDUST pipeline are available from ^84^ and can also be downloaded from https://github.com/papouinlab

## Methods

### Mice

All experiments were conducted in accordance with the National Institute of Health Guide for the Care and Use of Laboratory Animals and were approved by the Institutional Animal Care and Use Committee of Washington University in St. Louis School of Medicine (IACUC #20180184, #21-0372, #24-0346). Both male and female animals were used. Mice were housed in groups of 2-5 siblings (with exceptions, see below) on a 12-12 light-dark cycle (9am ON, 9pm OFF, or 11am ON, 11pm OFF) with *ad libitum* access to food and water. α7nAChR^flox/flox^ mice (Strain B6(Cg)-Chrna7^tm1.1Ehs^/YakelJ; JAX#026965), Glast-CreER^T2^ mice (Strain STOCK Tg(Slc1a3-cre/ERT)1Nat/J; JAX#012586), Camk2a-CreER^T2^ mice (Strain B6;129S6-Tg(Camk2a-cre/ERT2)1Aibs/J; JAX#012362), Gad2-CreER^T2^ mice (Strain STOCK Gad2^tm1(cre/ERT2)Zjh^/J; JAX#010702), Ai9 mice (Strain B6.Cg-Gt(ROSA)26Sor^tm9(CAG-tdTomato)Hze/J^; JAX#007909) were obtained from the Jackson Laboratory and bred in our facility.

Pimers used for genotyping were as follows:

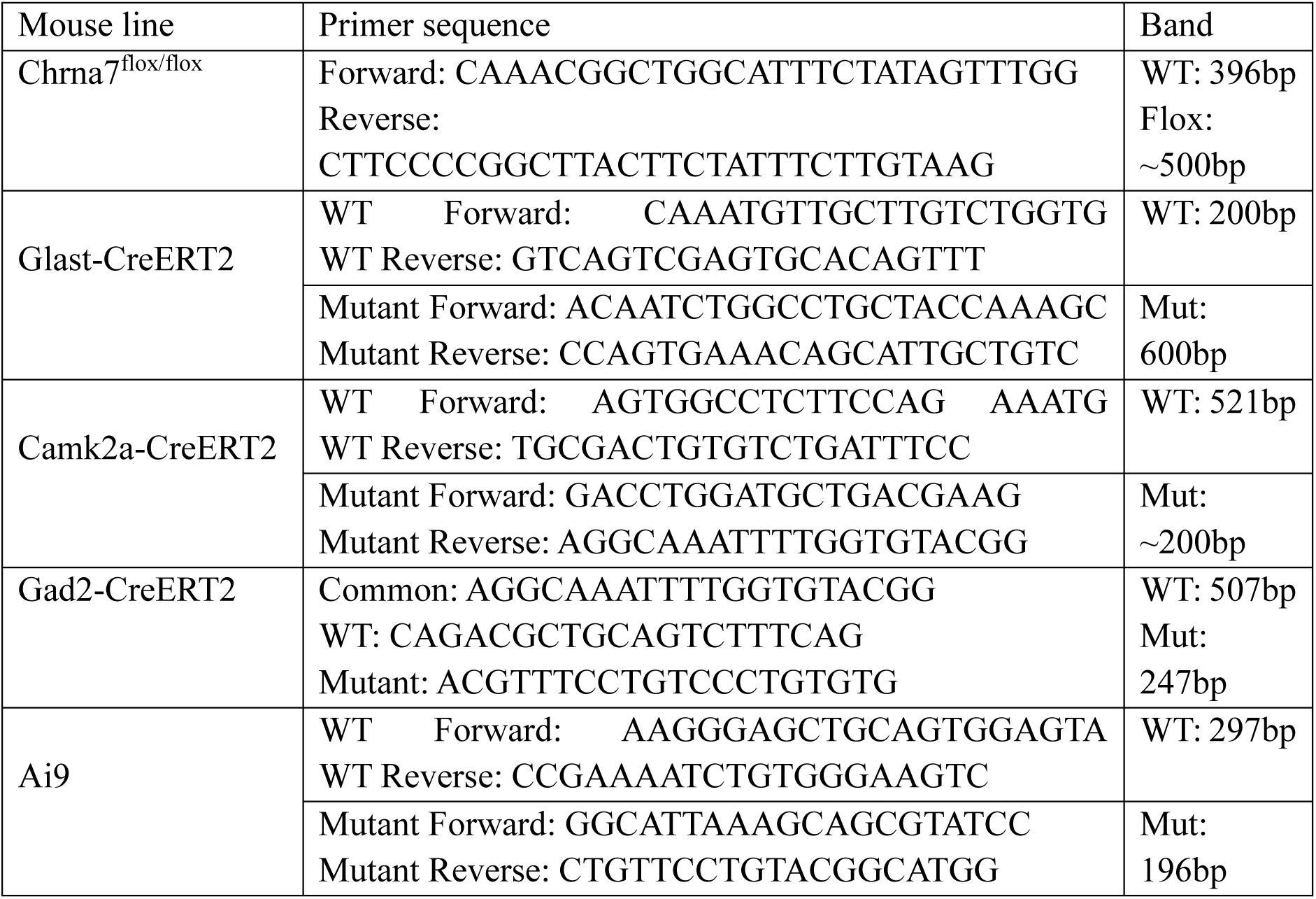

Astrocyte-specific α7nAChR KO mice, excitatory neuron-specific α7nAChR KO mice, and inhibitory neuron-specific α7nAChR KO were generated by crossing Chrna7^flox/flox^ mice with Glast-CreERT2 mice, Camk2a-CreERT2 mice and Gad2-CreERT2 respectively. Control littermates consisted of Chrna7^+/+^, Cre^+^; Chrna7^flox/flox^, Cre^-^ and Chrna7^+/flox^, Cre^-^ mice, treated with tamoxifen. Triple transgenic reporter mice (Chrna7^flox/flox^, Ai9^flox/-^, Cre^-^) were generated by crossing Ai9 mice with the conditional α7nAChR KO mice. Tamoxifen (Sigma T5648, 75mg/kg in corn oil for eN-α7^KO^ and A-α7^KO^, 150mg/kg for iN-α7^KO^) was given by i.p. injections for 5 consecutive days starting at P54 on average to induce recombination. Mice were used for behavior 5-6 weeks after tamoxifen treatment and for validation experiments no earlier than 4 weeks after tamoxifen treatment.

### Behavioral tests

The elevated plus maze, Y maze, novel object recognition task, open field, three chamber test and fear conditioning assays were conducted in an in-lab behavioral suit as described in Manno et al. 2020^85^, immediately adjacent to the mouse housing facility. Environmental control included restricted access, avoidance of extraneous scents and movements, and full control of light scenery. The testing space consisted of three sound-proof white walls with large and distinct dark visual cues, and was separated from the operator area by a black thick curtain. The battery of behavioral assays was administered longitudinally with the following sequence, with minor deviations when needed: elevated plus maze, Y maze, novel object recognition, three chamber test, ultrasonic vocalization, fear conditioning or pre-pulse inhibition, nesting, and sucrose preference test. EEG/EMG recordings were carried out on a separate cohort of animals.

Male and female mice were 12–15-week-old when used for behavioral testing. To minimize cage effects, tamoxifen-treated littermates with the following genotypes were used as controls: Chrna7^+/+^, Cre^+^; Chrna7^flox/flox^, Cre^-^ and Chrna7^+/flox^, Cre^-^. Behavioral tests were performed at the onset of the light-phase (Zeitgeber time 0, (ZT0), i.e. 9am or 11am) or at ZT6 (3pm or 5pm) for a maximum of 2 hours, depending on the assay. Mice were acclimated to the room for 15 minutes before testing and immediately returned to their home-cage and vivarium room thereafter. Behavior tracking and analysis were conducted using a SMART3.0 software (Panlab, SMART 3.0) coupled to a NIR camera tracking system. Triwise setting in Smart3.0 allowed for tracking of the nose, center of mass or entire body of the animals as desired. Testing was conducted under 230 Lux unless otherwise stated. Experimenters were blind to the genotype of experimental mice during both data collection and analyses, unless otherwise stated and justified (see below). Treatment with EVP-6124 or saline was conducted by a separate experimenter not in charge of behavioral testing. Mazes and objects (when applicable) were thoroughly cleaned before and after each trial with 70% ethanol and then wiped dry with a clean wipe. Mazes were cleaned weekly with a Quatricide disinfectant.

#### Elevated plus maze (EPM)

The elevated plus maze consisted of two open arms and two closed arms with tall walls as described in Manno, 2020 ^85^. All surfaces were Sintra 19-mm-thick PVC foam board (grey or black). Environmental brightness was set at 1060 lux to induce threat-driven avoidance behavior. Mice were placed at the center intersection facing one of the closed arms and allowed to freely explore for 5 minutes for a single session. Mouse detection and tracking started with a short 5 sec delay. Open arm exploration occurred when all 4 paws passed the threshold of the open arm (nose, center of gravity and tail base). Open arm poking consisted of partial arm entries (nose or nose + center of mass), during which animals typically stretched their body in the open arm while maintaining their hind paws within the center interaction zone of the maze.

#### Y maze (YM)

The Y maze consisted of three arms with tall grey walls made of Sintra 19-mm-thick PVC foam board as described in Manno et al^85^. Each wall featured distinct visual cues. Mice were placed at the arm intersection at the start and allowed 8 min of free exploration. Four zones were delineated on the Smart3.0 interface; one for each arm and one for the center triangle. An arm entry was defined as a full boundary crossing by the entire body of the animal (nose, center of mass and tail base). An alternation was defined as a rolling sequence of triplets of distinct arms exploration (e.g., A > B > C). The percentage of alternations were calculated as [(Number of alternations) / (total number of arm entries -2)] x 100%. With our setup, chance level was at 33% alternations.

#### Novel object recognition (NOR)

The novel object recognition task was conducted in a quadruple open field, each consisting of a square enclosure made of grey Sintra 19-mm-thick PVC foam board, allowing up to four mice to be run simultaneously. A three-day protocol was used, starting with an acclimation phase (day 1) consisting of 10-minute exploration of the empty arena. Training occurred 24 hours later (day 2) and consisted of two 10-minute sessions of exploration with two identical objects (30-min inter-trial interval). Objects were positioned in consistent locations via magnets underneath the apparatus. During the 30-minute inter-session interval mice were returned to their cage but remained in the room while the objects and arenas were cleaned. The probe phases (day 3) consisted of one 10-minute exploration session with one familiar object from the training day replaced with a novel never-encountered object. Two sets of objects were used with identical dimensions but differences in texture, patterns, material, color and shape (see Manno et al^85^). Objects were randomly assigned on the training day. Novel object placement at the preferred or non-preferred position (i.e. most or least explored on the training day) was consistent within cage but randomized across cages. Mouse nose within 2.5 cm of the object, directed at the object, was used as a criterion for object exploration. The discrimination index (DI) was calculated as [(exploration time of novel object – exploration time of familiar object) / (total exploration time)]. The same formula was used to calculate the novel object place preference in the acquisition phase. For EVP-6124 experiments the first 5 mins of probe exploration were used to calculate the discrimination index. Mice that only explored one of the objects or explored objects for less than 10 seconds combined during the combined two sessions of acquisition were excluded from the analysis. Pilot examinations indeed indicated that this is not sufficient for wild-type mice to build an object preference on the probe day. Mice that explored less than 2 seconds on the probe day were also excluded because this prevented accurate measurements of the discrimination index. In total, 19 out of 271 mice were excluded from analysis using these criteria (7%).

#### Three-chamber test (TCT)

The three-chamber test was run in a transparent Plexiglas arena consisting of three chambers separated by two partitions, each with a door operated by a remote pully system. Mice were initially placed in the middle chamber. At the start of the test, the side doors were opened, allowing the mice to freely explore all three chambers. The assay consisted of three consecutive phases of 10 minutes each. In the first phase, mice were allowed to explore all three chambers, with an empty upside-down wire cup secured in each of the side chambers. At the end of the first phase, mice were temporarily confirmed to the middle chamber while a stimulus mouse was placed in one of the cups and an object in the other. Doors were lifted again and the test mouse was free to explore all chambers for 10 min (phase 2). Proceeding identically, the object was replaced with a second stimulus mouse at the start of phase 3. Non-littermate age- and sex-matched mice were used as stimulus mice. The two stimulus mice were from distinct cages. The wire cups allow contact (and scent) between stimulus and test mice, but no biting or other aggressive behavior. No aggressive behavior was observed with any of the mice tested. The sociability index was measured as [time interacting with mouse / time exploring the mouse and object] during phase 2. The social preference index was measured as [time interacting with new mouse / time exploring the two mice] during phase 3. Three mice (out of 225) had to be excluded due to climbing on wired cups throughout the 3 phases.

#### Fear conditioning (FC)

The fear conditioning test was conducted as described in Papouin et al. 2017^40^ with minor modifications. Briefly, up to four mice were tested simultaneously in four rat conditioning chambers (Coulbourn Instruments, H13-15) placed inside sound-attenuating isolation cabinets. Two contexts were used that differed in scent (lemon vs almond), visual cues (original enclosure vs checkered pattern), texture (metal wall vs plastic), shape (square vs round), size (large vs medium) and brightness (bright vs dim). The grid floor through which shocks were delivered was identical in both contexts. We used a four-day protocol consisting of acclimation, fear leaning (shocks), fear retrieval, and test of context specificity. The acclimation and context-specificity tests were carried out in context B, while fear learning and retrieval were conducted in context A. The fear learning consisted of three 2-second shocks (0.7 mA) delivered 1min apart without auditory cue pairing, after 4-minutes of no-shock exploration. The habituation, retrieval, and context specificity tests consisted of 4 minutes of free exploration without shocks. Freezing was defined as the total absence of locomotion and body movement other than breathing for at least 1 second.

#### Ultrasonic vocalization (USV)

The ultrasonic vocalization test was carried out in a sound-attenuating isolation chamber equipped with an Avisoft UltraSoundGate CM16 microphone connected to an ultrasound recording amplifier (Avisoft UltraSoundGate 116H 1.1 #1). USVs between pairs of male and female mice of the same genotype were recorded with Avisoft Recorder software (gain = 4 dB, 16 bits, sampling rate = 250 kHz) over a 10-minute session. The estrous cycle of female mice was tracked daily for a week ahead of testing via vaginal smears^86^. Females in estrus or proestrus stages were used for testing. To promote social interactions and vocalizations, the male mouse was first habituated to the chamber with bedding collected from the female’s home cage for 10 min before the female mouse was introduced into the chamber. USVs were captured and analyzed from the sonograms using a customized pipeline as previously described^52,87^. Specifically, jump calls are defined as calls that contain a discontinuous jump in frequency of at least 10 kHz. Chunks refer to a sequence of calls separated by pauses of less than 0.5 seconds. Tuples refer to bouts containing more than 1 call. Singlets refer to bouts containing a single call.

#### Nesting

Nesting behavior was assessed in the animals’ home cage. For this assay, mice were singly housed and provided with 5 g of pressed cotton squares. Cages were left undisturbed. Nest building was assessed 4-6 hours later, and again at 24 hours. Nest scoring was adapted from the rating scale described by Deacon R.2006^88^ with minor modifications. Specifically, a score of 0 was given when cotton squares were left untouched. Score 1: Cotton squares nearly intact. Score 2: Cotton squares were shredded but only partially (50%-90% intact) and not assembled into a nest. Score 3: The squares were mostly shredded (less than 50% intact) and the shredded material was assembled into a partial nest, or >90% of the squares were shredded but not assembled into a nest. Score 4: The squares were entirely shredded and assembled into a clear nesting area. Score 5: A finished nest has been formed from fully shredded bedding material (usually into an igloo shape with an entry hole). To ensure consistency, all nest scoring was performed by a single experimenter.

#### Sucrose preference test (SPT)

The sucrose preference test immediately followed the nesting behavior assay and was carried out in the home cage. Mice were singly housed and given two graduated bottles containing water. Liquid consumption was measured daily by weighing the bottles. Mice were allowed to habituate to the two bottles for 3 days before one of the bottles was replaced with 1% sucrose solution. Consumption from both bottles was recorded for an additional 4 days. To account for side preference, the positions of the bottles were switched every day after weighing. Sucrose preference (%) was calculated by the [(consumption of sucrose – consumption of water) / total volume consumed] × 100.

#### Acoustic startle response and pre-pulse inhibition (PPI)

Whole-body startle response to an auditory stimulus pulses (40 ms broadband burst) and pre-pulse inhibition of the startle response PPI trials by preceding sub-threshold pre-pulse were measured concurrently in the same mice using the Kinder Scientific Startle Monitor system (Poway, CA). Test sessions began with a 5 min acclimation period in a 65 dB white noise background. A block of five consecutive startle trials was presented initially to habituate and stabilize the startle response in each mouse. 12 types of startle trials (70, 75, 80, 85, 90, 95, 100, 105, 110, 115, and 120 dB, in addition to no stim, each delivered 3 times) were interspersed with 4 blocks of PPI trials consisting of a 69, 73, 33 or 81 dB pre-pulse, 100ms before the 120dB startle pulse (8 trials for each PPI block). Inter-trial intervals were semi-random (5-25 seconds). Force readings (in newtons) following startle stimulus onset were averaged to obtain an animal’s startle amplitude. PPI was measured as %PPI= 100-(100*(startle amplitude to pulse alone – startle amplitude to prepulse + startle pulse)/startle amplitude to pulse alone). Startle responses at each dB level (3-4 trials per dB level) were averaged.

#### EEG/EMG recording

Electroencephalogram (EEG) cranial screws (Pinnacle Technology) and an EMG neck wire were implanted stereotaxically under isoflurane anesthesia on 6-8-week-old male C57BL/6J mice (Jackson Laboratory, 000664) as in Papouin et al., 2017^40^. Briefly, the surface of the skull was exposed and four insulated wire electrodes were placed and screwed as follows: Two extradural cortical electrodes were inserted bilaterally in the frontal areas and the other two were inserted bilaterally in the parietal areas. Two insulated wire electrodes were inserted bilaterally into the nuchal muscle for electromyogram (EMG) recordings. Electrodes connected to a micro-connector (Pinnacle technology) were secured at the surface of the skull with dental acrylic. Mice received buprenorphine (0.08 mg/kg) and saline i.p. and allowed 5 days of recovery. Lightweight recording cables (Pinnacle Technology) were connected to the head implants and mice were placed in Pinnacle Plexiglas cages containing water and food ad libitum for acclimation for 2-3 days in a sound-proof room. EEG and EMG signals were amplified and bandpass filtered at 0.5–100 Hz and 10–100 Hz, respectively, using a 15 LT Bipolar Physiodata amplifier system (Grass Technologies) and sampled at 400 Hz with a MP150 data acquisition system (BIOPAC Systems). The system was equipped with infra-red cameras to monitor animal behavior and activity throughout the recording. EEG and EMG waveforms were then analyzed as in Kohtoh et al. (2008) using SleepSign for Animal software (Kissei Comtec) with minor changes. Briefly, waveforms were analyzed by 10 s epochs (1024 FFT points). Each Epoch was divided into 5 regions that were FFT analyzed using a Hanning Window and averaged. Three frequency bands were calculated: δ (0.75-4Hz), θ (4-8Hz) and α (8-12Hz) and used to determine vigilance states using the following criteria with the indicated order of priority: 1-Clear locomotion or EEG integral > 3-5 μV/sec = active wakefulness; 2-EEG integral < 3-5 μV/sec and EEG δ power > 250-500 μV2 = Non-REM sleep; 3- EEG integral < 3-5 μV/sec and EEG θ/(δ+ θ) ratio and/or EEG α /(δ+ α) > 45% = Non-REM sleep; 4- otherwise = quiet wakefulness.

#### Locomotion in open field (OF)

The open field assay of locomotion was conducted during the acclimation phase of the NOR. Briefly, mice were allowed to freely explore an empty open field for 10 min under the dim lighting (230 lux). Distance travel was measured using Smart3.0.

Body weight: Animal body weight was measured at P90-100 before the start of behavioral tests.

### Z-scoring of behavior performance

To compare performance across lines, a compounded behavioral z-score was obtained for each behavior, calculated as the average z-score across relevant parameter, as follows. YM: spontaneous alternation (%), NOR: discrimination index, phase 3; TCT: phase 2 and phase 3 social discrimination indexes; FC: percentage of time spent freezing during fear learning and percentage of time spent freezing during fear retrieval; Startle: average responses to the high intensities (105 dB, 110 dB, 115 dB and 120 dB); PPI: pre-pulse inhibition at the 69dB, 73dB, 77dB, 81dB pre-pulse; USVs: total call number and total number of jump calls; EPM: time spent in the open arms and time spent poking; Nesting: score at 4 hour and score at 24 hour; SPT: total sucrose preference; OF: total distance travelled; Weight; Sleep: power spectrum across different stages, time spent in each stage in light, and time spent in each stage in dark.

### D-serine Supplementation

D-serine supplementation was administered through drinking water to minimize stress. D-serine (Sigma S4250, 1.5 to 2.4mg/mL) was directly dissolved in the drinking water and provided ad libitum for 7 days before behavioral testing and throughout the subsequent testing period. No other source of water was provided during this time. This protocol resulted in a daily dose of approximately 150-400 mg/kg D-serine per body weight considering an average weight of 30 g and drinking volume of 5 ml/day (calculated during the two-bottle assay)

### EVP-6124 treatment

EVP-6124 (EVP, MedChem Express, HY-15430A, 5 mg) was dissolved in DMSO into 10mg/mL or 50mg/mL stock solutions as needed. The appropriate amount of EVP-6124 in DMSO was diluted with saline on the day of injections at the desired final concentration. Dose response curves (0.1 mg/mL, 0.3 mg/mL, 0.6 mg/mL, 1 mg/mL) were obtained in naïve C57BL/6J male mice (Jackson Laboratory, 000664). Littermate controls received vehicle (saline with matched amount of DMSO). Both male and female A-α7^KO^ and A-α7^control^ mice that had not undergone other behavioral testing were used for EVP-6124 experiments. The experimenter was blind to the treatment and genotype. These experiments were conducted at ZT6. All animals received EVP-6124 30 min (males) or 60 min (females) prior to the start of the second phase (identical objects). Mice were returned to the home cage after injection until used for testing.

### Pharmacokinetic studies

Pharmacokinetic studies of EVP-6124 were performed by Paraza Pharma, Inc. (Montreal, Canada), study ID: C050-012/PK-2020-156. Briefly, all animals were weighed, and the dosing formulation volumes were calculated accordingly. EVP-6124 was administered intraperitoneally (i.p.) to all animals (with the exception of pre-dose animals). At selected time points (Predose, 0.5, 1, 2, 4, 7 and 24 hours post-dose), the animals were anesthetized with isoflurane and transcardially perfused with phosphate buffered saline (PBS, pH 7.4). The brains were then harvested and homogenized by polytron 1:4 (w/v) in 25% isopropanol in water. EVP-6124 was then detected and quantified on a Quantis LC/MS system. Briefly, 50 μL of brain homogenate sample were mixed with 50 μL of spiking solution and 150 μL of internal standard working solution (0.1 μM Glyburide/Labetalol in Methanol/ACN), vortexed, and centrifuged (10,000g for 10 min at 4°C). 200 μL of supernatant was transferred to a 2mL deep well plate and evaporated to dryness at 60°C, then reconstituted with 200μL of reconstitution solution and vortexed before being injected on LC-MS/MS (20µL). Samples were run on a Xbridge BEH C18 2.1 x 30 mm, 2.5 μm column at 50°C. Non-compartmental analysis (NCA) was performed on the composite pharmacokinetic profiles (plasma and brain) using WinNonlin 8.1.

### Stereotaxic microinjections

Mice were anesthetized with 5% vol/vol isoflurane and maintained under 1%-2.5% vol/vol of continuous isoflurane (vaporizer RWD) and placed on a surgical stereotaxic frame (KOPF instruments). 1mg/kg Buprenorphine Extended Release (Wedgewood Connect) was administered subcutaneously before surgery. After hair removal and disinfection with 70% ethanol and 10% povidone iodine at the surgical site, an incision was made, followed by bilateral craniotomy using a high-speed rotary micromotor (Foredom). All injections in the present study were targeted to the hippocampal CA1 region, with coordinates of -2AP, ±1.5 ML, -1.4 DV (in mm) from Bregma. Adeno-associated viruses (AAVs) were microinjected at a rate of 150 nL/min with a 2 µL syringe (Hamilton 7002, bevel tip, 30 gauge) controlled by a syringe pump (KD Scientific). After injection, the syringe was left in place for at least 8 minutes before withdrawing. The surgical opening was closed by 6-0 nylon sutures. Mice were closely monitored for 4 days and used no sooner than 3 weeks post-surgery. Viruses used in the present study were as follow: 1 µL/site of 9:1 co-injection of AAV5-*gfaABC1D*::lckGCaMP6f (Addgene #52924-AAV5, 2.5 x 10^13^ GC/ml) and AAV5-*gfaABC1D*::tdTomato (Addgene #44332-AAV5, 2.2 x 10^13^ GC/ml).

### Preparation of acute brain slices

Acute hippocampal coronal slices (350µm) were prepared from 12-15-week-old mice as described previously^40,43,89^ and used for calcium imaging or electrophysiology. Briefly, mice were deeply anesthetized with isoflurane and decapitated. The brain was quickly removed from the skull and sectioned on a Leica VT1200s vibratome to obtain 350 µm slices. Slicing artificial cerebrospinal fluid (aCSF) composition was as follows: 125 mM NaCl, 3.2 mM KCl, 1.25 mM NaH_2_PO_4_, 26 mM NaHCO_3_, 1 mM CaCl_2_, 2 mM MgCl_2_, and 10 mM glucose saturated with 95% O_2_/5% CO_2_ (pH 7.3, osmolarity 290-300 mOsm.L^-1^). The acute brain slices were then maintained at 33℃ for 30 minutes and then at room temperature for 45 minutes before being used for experiments in normal aCSF: 125 mM NaCl, 3.2 mM KCl, 1.25 mM NaH_2_PO_4_, 26 mM NaHCO_3_, 2 mM CaCl_2_, 1.3 mM MgCl_2_, and 10 mM glucose saturated with 95% O_2_/5% CO_2_ (pH 7.3, osmolarity 290-300 mOsm.L^-1^).

### Field recordings

Acute brain slices were placed into the chamber of a Scientifica SliceScope Pro 6000 system where they were perfused with aCSF saturated with 95%O2/5%CO2 at a flow rate of ∼1mL/min. The aCSF was maintained at 33°C (TC344C Dual Channel Temperature Controller, Warner Instruments). Shaffer collaterals were electrically stimulated at 0.05 Hz with paired stimulations (0.1 ms pulse, 200 ms apart) using a concentric tungsten electrode placed in the *stratum radiatum*. Evoked field excitatory post-synaptic potentials (fEPSPs) were recorded using a glass electrode (2-4 MΩ) filled with recording aCSF placed in the *stratum radiatum* >200 µm away from the stimulus electrode. For long-term potentiation (LTP) experiments, stimulation (<150 µA, 100 µs) was set to ∼70% of the intensity that triggered population spikes. A 20-minute baseline was recorded before three trains of high-frequency stimulation (HFS, 100 Hz, 1 s) were delivered, 20s apart. The slope of fEPSPs was monitored for the subsequent 50 minutes. All experiments were performed at 33 ℃ in the presence of GABA_A_ receptor blocker picrotoxin (Fisherscientific 11281G, 50 µM). For D-serine rescue experiments, D-serine (Sigma S4250, 20 µM) was added to aCSF 15 minutes before HFS. Data were acquired with a Multiclamp 700B amplifier (Molecular Devices) through a Digidata 1440A, sampled at 20 kHz, filtered at 10 kHz, and analyzed using pClamp11.0 software (Molecular Devices). LTP potentiation was determined by measuring the slope of fEPSP traces, averaged from 30 weeps obtained 41-50 min post HFS stimulation and compared to the 10 min baseline that immediately preceded HFS delivery.

### Two-photon calcium imaging

GCaMP-based fluorescent recordings of calcium activity in CA1 *stratum radiatum* astrocytes were performed in acute hippocampal slices prepared as above. Brain slices were maintained in recording aCSF saturated with 95% O_2_/5% CO_2_ with a perfusion rate of 2.5 mL/min during recording. Recordings were obtained from a Nikon A1R or Bruker Ultima 2pPlus, a two-photon laser scanning microscope (excitation wavelength 920 nm, laser power 37.5 mW, 515/530 filter) with a 25X water-immersion objective (Nikon objective 25X, 1.10 NA in both cases). Astrocyte calcium signals were recorded at 40-60 µm depth at 1 Hz with 1.6X digital zoom. All recordings were obtained in the presence of tetrodotoxin (1 µM) to prevent the influence of neuronal activity. Drugs used were as follows: PNU-282987 (Tocris 2303, 500 nM), MLA (Methyllycaconitine citrate, Tocris 1029, 100-500 nM). Recording aCSF composition was: 125 mM NaCl, 3.2 mM KCl, 1.25 mM NaH_2_PO_4_, 26 mM NaHCO_3_, 2 mM CaCl_2_, 1.5 mM MgCl_2_, and 10 mM glucose saturated with 95% O_2_/5% CO_2_ (pH 7.3, osmolarity 290-300 mOsm.L^-1^).

### Calcium analysis

Astrocyte calcium recordings were analyzed with the STARDUST pipeline^84^.

Briefly, a map of regions of activity (ROA) was obtained from the temporal projection of active pixels identified by the AQuA2 software ^90^. This map of ROAs was then overlaid onto the raw movie to extract ROA-based time-series data. Where required, a cell map was obtained by hand-drawing the cell outlines based on stable fluorescence markers during recording. The ROA map, ROA-based time-series data and cell map were fed into STARDUST to extract signal features (amplitude, frequency, kinetics) from normalized signals (ΔF/F_0_) and obtain cell-based ROA data (cell-based signal amplitude, frequency, kinetics, number of ROAs per cell, size of ROAs per cell) by assigning ROAs to individual astrocyte. Spontaneous transients were analyzed from 10 minutes of recording. The drug epoch was defined as 100 - 300 seconds post drug application for PNU-282987.

### Immunohistochemistry

Mice were deeply anesthetized with isoflurane and transcardially perfused with 20-30 ml PBS followed by 20-30 ml 4% paraformaldehyde (PFA, Thermo Scientific). Brains were gently removed from the skull and post-fixed in 4% PFA for 24 hours at 4℃, followed by dehydration in 30% sucrose (w/v) in PBS for 2 days at 4 ℃. 50 µm free-floating sections were obtained with a vibratome (Leica VT1000) in cold PBS and stored at 4℃ in PBS with 0.02% sodium azide (Sigma-Aldrich, RTC400068) for long-term storage. For immunofluorescence staining, brain slices were first permeabilized in 0.5% Triton X-100 dissolved in PBS (PBST) for 10 minutes, followed by blocking with 5% normal goat serum (NGS, Abcam, AB 7481) in 0.5% PBST for 30 minutes. Sections were then incubated with primary antibodies diluted with 1% NGS in 0.5% PBST at 4℃ overnight with an orbital shaker. The next day, sections were washed with PBS for 5 minutes 3 times and incubated with secondary antibodies diluted in 1% NGS in 0.5% PBST for 2 hours at room temperature protected from light. Sections were then washed three times with PBS. Second wash with DAPI (Sigma-Aldrich, MBD001, 1:1000) was 10 minutes, while the other two washes were 5 minutes. Finally, sections were mounted on microscope slides (Fisher Scientific, 1255015) with mounting medium (Vector Laboratories, H-1400) and glass microscope cover slips (Fisher Scientific, 12541024). Slides were left drying in the dark at room temperature for a few hours before sealed with nail polish. Primary antibodies used in the present study were as follow: guinea-pig anti-NeuN (1:500, Millipore Sigma, ABN 90), rabbit anti-GFAP (1: 1000, Abcam, ab7260), rabbit anti-S100β (1:1000, Abcam, ab52642), rabbit anti-Sox9(1:500, Abcam, ERP14335-78), mouse anti-Gad1(1:500, Abcam, K87) Secondary antibodies were goat-anti-rabbit Alexa Fluor 488 (1:1000 ThermoFisher, A-11034) and goat-anti-guinea pig Alexa Fluor 633 (1:1000, ThermoFisher, A-21105), goat-anti-mouse Alexa Fluor 633 (1:1000, ThermoFisher, A-21052).

### Confocal imaging and image analysis

Confocal images were acquired on a Zeiss LSM 980 confocal microscope with a Plan-Apochromat 20×/0.8 air immersion objective (Zeiss, 440640-9903-000) using 405, 488, 561, and 633 nm laser lines. Z-stack images were acquired at steps of 1.0 μm apart imaging an area of 424.26 × 424.26 μm at 1024 × 1024 pixels resolution.

Cell counting and colocalization analysis were performed as described in Lefton et al. 2025^43^ with minor modifications. Briefly, GFAP and Sox9 were used to identify astrocytes in regions of CA1 *stratum radiatum* and dentate gyrus; S100β and Sox9 were used to identify astrocytes in regions of the cortex and amygdala. NeuN was used to identify neurons for all regions. Due to the complex structure of astrocytes, all cell counting and colocalization analysis were done manually on stacks of images without z-projections. Only astrocytes with cell bodies appearing in the stacks were counted. For experiments with Ai9 reporter mice, specificity was quantified as the percentage of tdTomato-positive cells that also expressed the desired cell marker(s). Efficiency was quantified as the percentage of cell-marker positive cells that also expressed tdTomato.

### DNA, RNA extraction, reverse transcription, RT-qPCR & PCR

Mice were deeply anesthetized with isoflurane and transcardially perfused with 20-30 mL PBS to remove blood. Brains were gently removed from the skull and quickly placed onto cold glass slides. 30 mg tissue was then taken from a ∼500 μm coronal hippocampal slice and maintained on ice. DNA and RNA extraction were performed with respective extraction kits according to the kit manuals (RNeasy Mini kit, Qiagen 74104; DNeasy Blood and Tissue kit, Qiagen 69504). RNA (1 ug per 20 µL reaction volume) was immediately converted to cDNA using EasyScript cDNA synthesis kit (v2.0, Lamda Biotech, G234). RT-qPCR was then performed to quantify KO efficiency from a transcripts level. All qPCR experiments were performed in triplicates using PowerUp SYBR green master mix (Thermo Fisher, A24742) on MicroAmp fast optical 96-well reaction plates (Applied Biosystems, 4346907) with QuantStudio6 (Applied Biosystems). A total of 40 cycles were run for all qPCR experiments. For qPCR quantification, target gene expression was quantified relative to *Actb* and normalized to control group samples based on analysis method described in Taylor et al. 2019^91^.

PCR of genomic DNA for were performed with Platinum II Hot-Start Green PCR Mater Mix (Invitrogen) and the following cycling conditions: 94°C for 2 min, 94°C for 15 s, 60°C for 15s, 68°C for 15s, repeat 2-4 for 35 cycles, and hold at 4° C. For *Chrna*7 KO allele detection, step 4 was extended to 38 s. Gel (1% agarose) was imaged with ChemiDoc MP Imaging System (Biorad, 5s exposure).

RT-qPCR primers were as follow: primers spanning *Chrna7* exon 4-5 (floxed exon): Forward 5’-ACATGTCTGAGTACCCCGGA-3’, Reverse 5’- CAAAGCGTTCATCTGCACTGT-3’ *Actb*: Forward 5’-GGCTGTATTCCCCTCCATCG-3’, Reverse 5’-CCAGTTGGTAACAATGCCATGT-3’ PCR primers for KO allele detection: Forward 5’-GAGGCTCAGGGTTTGAGGCA -3’, Reverse 5’- GGTCACTGAGCAGTGGTGGACAG -3’ *Tbp*: Forward 5’-CCTTGTACCCTTCACCAATGAC-3’, Reverse 5’- ACAGCCAAGATTCA CGGTAGA -3

### In vivo micro-dialysis and HPLC

This procedure was performed as described in Papouin et al. 2017^40^ with minor modifications. Briefly, mice were unilateral implanted with a guide cannula immediately above the hippocampus via stereotaxic surgery at P90-150. Coordinates were -2AP, -1.5 ML, -1DV (in mm) relative to Bregma. The guide cannula (CMA P000138) was secured with dental cement and two anchor cranial screws. Mice recovered for 5-7 days before sampling. Cannula implantation was assessed at the end of the experiment with brain sectioning. Micro-dialysate samples from off-targeted mice were excluded from the subsequent analysis. Micro-dialysis was carried out in a custom-made semi-transparent acrylic chamber. Mice were acclimated to the chamber and room for at least 1-2 days. On the day of sampling, probes (CMA P000082) were inserted through the guide cannula at ZT0. ACSF (Harvard Apparatus, 597316) was perfused through the probe at a rate of 0.5ul/minute for 6 hr. Dialysis samples were collected every hour on a fraction collector (BAS HoneyComb MF-9096) at 4℃ and the frozen at -20℃.

D-serine detection was conducted using chiral high performance liquid chromatography (HPLC) on a (Shimadzu Prominence-i LC2030C plus) using an Accucore C18 column (Thermo Fisher, 17126-152130). Aqueous phase (Buffer A, 0.05 mol/L disodium hydrogen phosphate, pH 6.5) and organic phase (Buffer B, 1:1 water and methanol with 0.05 mol/L disodium hydrogen phosphate, pH 6.5) were perfused at a rate of 0.6 mL/minute in a step protocol: 0-5.5 min, 6% mobile phase B; 5.5-8 min, 6% mobile phase B to 10% mobile phase B; 8-12 min, 10% mobile phase B; 12-16 min, 10% mobile phase B to 40% mobile phase B; 16–20 min, 40% mobile phase B; 20-25 min, 40% mobile phase B to 6% mobile phase B; 25-35 min, 6% mobile phase B. Samples were derivatized using o-Phthaldialdehyde (Sigma, P0657) and N-Acetyl-L-Cysteine (Sigma, A7250) for 5 minutes. After derivatization, 50 µL acetic acid (1 mol/L) was added to stop the reaction. 5 µl of derivatized sample was injected and ran through the column in duplicates. Standards were prepared from D- and L-serine (Sigma, S4250 and S4500) in ultrapure water at a ratio of 1:10. The area under the D-serine peak was used to calculate the sample concentration relative to standards.

### Statistical analysis

Statistical analysis was conducted using Graphpad Prism 10 software (v.10.4.2) and Python (v. 3.11.13). Prior to analyses, Shapiro-Wilk test was performed to determine normality. Student’s *t*-test (including multiple Student’s *t*-tests), Mann-Whitney test (including multiple Mann-Whitney tests), paired Student’s *t*-test, Wilcoxon test, one-way ANOVA and Tukey’s multiple comparisons test, two-way ANOVA and Sidak’s multiple comparisons test were then performed on where appropriate. Permutation test was performed on astrocyte calcium data (permutations set at 10,000). Chi-square test was performed to assess calcium ROA type distribution. Sex × genotype interactions are summarized in Table 1. Statistical data on all figures are reported collapsed for sex. All statistical data can be found in Table 2.

